# Trim69 is a microtubule regulator that acts as a pantropic viral inhibitor

**DOI:** 10.1101/2022.06.20.496826

**Authors:** Yuxin Song, Xuan-Nhi Nguyen, Anuj Kumar, Claire da Silva, Léa Picard, Lucie Etienne, Andrea Cimarelli

## Abstract

To identify novel cellular modulators of HIV-1 infection in IFN-stimulated myeloid cells, we have carried out a screen that combines functional and evolutionary analyses in THP-1-PMA cells that led us to the Tripartite Motif Protein 69 (Trim69), a poorly studied member of the Trim family of innate immunity regulators. Trim69 inhibits HIV-1, primate lentiviruses and the negative and positive-strand RNA viruses VSV and SARS-CoV2, overall indicating it is a broad-spectrum antiviral factor. Trim69 binds directly to microtubules and its antiviral activity is intimately linked to its ability to promote the accumulation of stable MTs, a specialized subset of microtubules. By analyzing the behavior of primary blood cells, we provide evidence that a program of MT stabilization is commonly observed in response to IFN-I in cells of the myeloid lineage and Trim69 is the key factor behind this program.

Overall, our study identifies Trim69 as the first antiviral innate defense factor that regulates the properties of microtubules to limit viral spread, highlighting the possibility that the cytoskeleton may be a novel unappreciated fighting ground in the host-pathogen interactions that underlie viral infections.

## INTRODUCTION

Like it is the case for all obligate intracellular parasites, viruses must compose with a complex cellular environment. Among the cellular components that play negative effects on virus replication are effectors of type I interferon responses (IFN-I), or interferon-sensitive genes (ISGs) [1]. While as part of a whole they participate to a generalized anti-pathogen state, individual ISGs can be either specific for one class of viruses, or can inhibit processes common to very diverse viruses, as for example the double-stranded RNA-activated protein kinase R (PKR), or the interferon-induced transmembrane proteins (IFITMs) that inhibits virus-to-cell membrane fusion [2–6]. In recent years, the search for cellular effectors directed against the HIV-1 retrovirus has drawn particular interest, spurring a number of genetic screens that are defining the complex cellular landscape in which HIV replication occurs (for example [7–12].

In search for novel ISGs that could interfere with HIV-1 infection, we have examined the weight of more than 400 ISGs during HIV-1 infection of IFN-stimulated macrophage-like cells (THP-1-PMA differentiated), that represent a cellular context particularly restrictive to infection [13]. Using a three-layer screen approach that combines functional and evolutionary analyses, we have identified the Tripartite Motif protein (Trim69) as a novel regulator of the early phases of the life cycle of HIV-1. Trim69 is a poorly studied member of the large Trim family that includes more than 80 members largely devoted to innate immunity regulation [14, 15]. In the past, Trim69 had been controversially linked to apoptosis and p53 signaling [16–19], but evidence of strong positive selection suggested this protein could be potentially involved in a host-pathogen genetic conflict [20]. Evidence that this could be the case came only in 2018, when Trim69 was reported to inhibit Dengue virus replication through the degradation of the viral non-structural protein 3 (NS3) [21]. While this finding could not be confirmed by a subsequent study [22], two studies identified Trim69 as an inhibitor of the Vesicular Stomatitis Virus (VSV) [22, 23]. The underlying mechanism of viral inhibition and more importantly the spectrum of viruses that can be targeted by this protein remain unclear.

In this study, we determine that in myeloid cells Trim69 is capable of inhibiting not only HIV-1, but also other primate lentiviruses, in addition to the negative-strand RNA virus VSV, and to the positive-strand RNA Coronavirus SARS-CoV2. These viruses are inhibited to varying degrees depending on the combination between virus and cell type, but overall, our results indicate that Trim69 is a novel antiviral factor endowed with a broad spectrum of inhibition. Trim69 inhibits the early phases of the above-mentioned viruses at entry and early post-entry phases and in particular reverse transcription, entry-primary transcription and RNA replication for HIV-1, VSV and SARS-Cov2, respectively. Antiviral inhibition is linked to the ability of Trim69 to induce the accumulation of stable microtubules, property that we described here is a previously unrecognized common cellular response to IFN stimulation. Overall, our results uncover Trim69 as a novel key modulator of microtubule dynamics during viral infection and further stress the importance that the control of the cytoskeleton network plays during the host-pathogen conflicts that underlie viral infection.

## RESULTS

### A three-layer genetic screen to identify novel mediators of the negative effects of IFN-I during the early phases of HIV-1 infection in macrophage-like cells

The early phases of HIV-1 infection are inhibited by type 1 interferons and more potently so in cells of the myeloid lineage [13]. To identify novel effectors of this antiviral response, we individually silenced >400 ISGs described in the *Interferome* database by lentiviral-mediated shRNA transduction in THP-1 cells that were split in two then differentiated into a macrophage-like state with PMA. We then examined the susceptibility of these cells to HIV-1 in the presence or absence of IFNα, using single-round of infection competent viruses coding a Firefly Luciferase reporter. IFNα modulators were defined as silenced genes that modulated the no IFNα/IFNα infectivity ratio (referred to here as the IFN defect) when normalized to the library average value (scheme of **Figure 1A**). To exclude modulators of IFNα signaling rather than of HIV-1 infection *per se,* a secondary screen was carried out on primary IFNα hits, by measuring the effects of gene silencing on the production of IP10, a well-described ISG, in THP-1-PMA cells stimulated with IFNα. A tertiary evolutionary analysis was then performed on remaining hits to prioritize them according to their evolutionary history and their potential roles in host-pathogen evolutionary genetic conflicts, using the DGINN pipeline (Detection of Genetic Innovation) [24].

**Figure 1.**
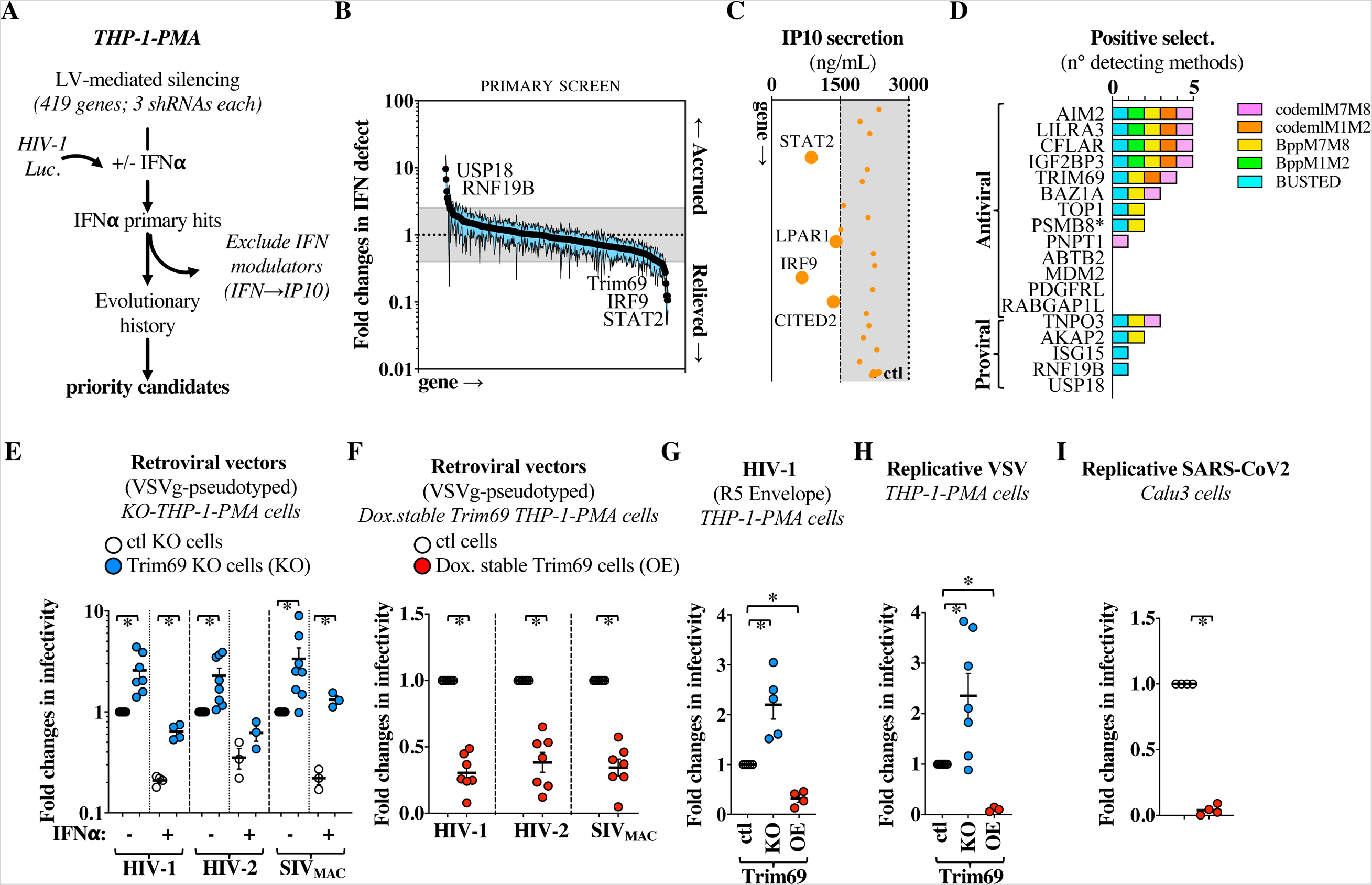
A three-layer genetic screen for IFN modulators of macrophage infection by HIV-1 highlights Trim69 as a broad viral inhibitor. A) Schematic approach used here. To identify IFN modulators of the early phases of HIV-1 infection in macrophages, 419 ISGs were individually silenced in THP-1 cells by lentiviral-mediated transfer. Cells were then divided in two, differentiated into a macrophages-like status with PMA, treated or not with 1.000U/mL of IFNα2 for twenty-four hours prior to viral challenge with a VSVg-pseudotyped HIV-1 virus coding for a Firefly Luciferase reporter. A secondary screen was carried out on primary hits to exclude genes that interfered with IFN signaling and a tertiary evolutionary screen was carried out on remaining candidate genes to focus on those most likely involved in an evolutionary-arms race. B) An IFN defect was calculated for each gene as a ratio between the luciferase activities obtained in the no IFN/IFN conditions that was normalized to the average value of the entire library. The graph presents AVG and SEM obtained. A threshold of 2.5 was chosen (outside the grey area) with values >2.5 that indicate accrued IFN defect and values <2.5 that highlight a relieved one. C) Cells silenced for the hits retrieved in B were stimulated with 1.000U/mL of IFNα2 and IP10 was measured by ELISA, one day later. Statistically significant differences in IP10 secretion following a one-way Anova test Dunnett’s multiple comparison test are represented with larger dots. D) DGINN-mediated positive selection analysis of hits retained after the secondary screen. The graph presents the number of methods that detect positive selection and proteins are separated according to the anti and pro-viral recategorization provided in the extended data Figure 1B. Of note, DGINN also identified a novel recombination event in PSMB8 and retrieved paralogues for LILRA3 and IGF2BP3 (extended data Figure 2C). E-G) THP-1 cells either ablated for Trim69 (KO) or overexpressing (OE) Trim69 under the control of a doxycycline promoter were challenged with the indicated single round of infection viruses expressing a GFP reporter at multiplicities of infection (MOIs) comprised between 2 and 4. The extent of infection was then measured two-three days later by flow cytometry. H-I) Replicative VSV and SARS-CoV2 viruses bearing respectively a GFP and mNeonGreen reporter were used to infect either THP-1-PMA cells or epithelial lung Calu-3 cells at MOIs comprised between 0.1 and 0.5. The extent of infection was measured either 16 hours or two days later by flow cytometry. All panels presented AVG and SEM of 3 to 8 independent experiments after normalization. Non-normalized values along with a WB analysis of Trim69 expression levels are presented in the extended data Figure 5. When indicated, IFNα was used at 1.000 U/mL for 24 hours prior to infection. *, indicates statistically significant differences between the relevant conditions, according to a two-tailed Student t test.

Under these conditions, the primary functional screen yielded 22 genes with significant impact on HIV-1 infection in the presence of IFNα (2.5-fold changes, **Figure 1B** and **extended data Figure 1** for individual values). Given that the IFN defect ratio can also be influenced by the susceptibility to infection of non-stimulated cells (due for instance to different basal levels of expression of the different genes), candidate genes were recategorized as anti- or pro-viral factors, according to the behavior of silenced THP-1-PMA cells during HIV-1 infection in the absence of IFN (**extended data Figure 1**). Given that silencing of AIM2, TOP1, IGF2BP3 and ABTB2 led to higher HIV-1 infection of unstimulated THP-1-PMA cells, these proteins were recategorized as antiviral factors. Conversely, silencing of AKAP2 and TNPO3, a well-known HIV cofactor [7, 25] that is not IFN-stimulated but that was introduced in our screen to serve as a sentinel gene, led to lower infection rates in unstimulated cells, so that these proteins were recategorized as pro-viral factors. The remaining genes did not exhibit significant changes in infectivity in unstimulated cells and were thus not recategorized. Overall, this analysis led to the identification of 5 pro- and 17 anti-viral modulators (**extended data Figure 1**) that were then analyzed for their ability to interfere with IP10 secretion upon IFNα stimulation (**Figure 1C**). Under these conditions, silencing of STAT2, IRF9 as well as of the CBP/p300-interacting transactivator with glutamic acid/aspartic acid-rich carboxyl-terminal domain 2 (CITED2) and the lysophosphatidic acid receptor 1 (LPAR1) significantly decreased IP10 secretion (**Figure 1C**). While the results obtained with STAT2 and IRF9 were expected [26], our results highlight a novel role for CITED2 and LPAR1 in the establishment of an IFN state. For the purpose of this study, the four above-mentioned genes were discarded from subsequent analyses. Most antiviral restriction factors are engaged into molecular evolutionary arms-races with pathogens [27, 28]. They therefore present signatures of these conflicts that can be identified by studying the evolution of their orthologs in host sequences [29]. To identify the candidate genes that present such genetic innovations during primate evolution, we screened the functionally retrieved hits with the DGINN pipeline [24], which automatically reconstructs multiple sequence alignments and phylogenies and that identifies events of gene duplication, recombination as well as marks of positive selection based on a coding sequence (**Figure 1D**). Automated or manually-retrieved sequences of primary hits were used as query for DGINN. We performed analyses in two steps: (1) phylogenetic analyses and (2) positive selection analyses that combine five methods from PAML Codeml, HYPHY BUSTED and bpp packages (see methods). We found five genes with evidence of strong positive selection in primates, detected by at least four methods (**Figure 1D** and **extended data Figure 2A, 2B and 2C**): Absent In Melanoma 2 (AIM2), the leukocyte immunoglobulin like receptor LILRA3, the CASP8- and FADD-like apoptosis regulator (CFLAR), the 6-methylated adenosine (m6A) reader IGF2BP3, and the tripartite motif protein 69 (Trim69), a poorly studied member of the TRIM family. Five additional genes exhibited some evidence of positive selection, detected by 2-3 methods: the Bromodomain Adjacent to Zinc Finger Domain 1A (BAZ1A), the DNA Topoisomerase I (TOP1), the A-Kinase Anchoring Protein 2 (AKAP2), the Proteasome subunit beta type-8 (PSMB8) and Transportin 3 (TNPO3) (**Figure 1D**).

### Trim69 is a novel broad inhibitor of viral infection

Among the top hits, we focused on Trim69, a prototypical member of the C-IV subfamily of Trims [30], which includes Trim5α which is a well-established antiviral factor against primate lentiviruses. Trim69 possesses a RING, B-box and Coiled-coil (CC) domains followed by a PRY-SPRY domain. Trim69 bears 48% and 44% identity with Trim67 and Trim52/Trim47 following a Blast analysis, while its SPRY domain is closer to Trim39 and Trim21 (46 and 45% identity, respectively, **extended data Figure 3A**). Comparison between the i*n silico* modeled SPRY domains of Trim69 and Trim5α indicates that Trim69 lacks the protruding variable loops (V1, V2 and V3) with which Trim5α contacts specifically retroviral capsids, suggesting an altogether different method of action (**extended data Figure 3B**).

Despite having being first identified as a spermatids-specific gene [17], Trim69 exhibits a heterogeneous pattern of expression in different cell lines and primary blood cell types tested (**extended data Figure 4A**). In THP-1-PMA cells, Trim69 is expressed already at basal levels and is extremely sensitive to IFN-I (α/β) stimulation, similarly to primary macrophages. To first validate the results obtained in our screen, we generated Trim69 knockout THP-1 cells (KO) by Crispr/Cas9 mediated gene deletion (**extended data Figure 4B** for cell viability) and then challenged macrophage-differentiated cells with HIV-1, HIV-2 or SIV_MAC_ lentiviruses in the presence or absence of IFNα. Under these conditions, removal of Trim69 increased the susceptibility of target cells to the three primate lentiviruses in the presence, but also in the absence of IFNα (from 2.25 to 3.35 on average), in line with its expression pattern (**Figure 1E**). Conversely and as expected for an antiviral factor, the susceptibility of THP-1 PMA cells stably expressing TRIM69 under the control of doxycycline (OE, **extended data Figure 4B** for cell viability) was decreased to the same extent during infection with these retroviruses (**Figure 1F**). Given that in this set of experiments viruses were pseudotyped with the pantropic envelope VSVg, Trim69 KO and OE cells were also challenged with an HIV-1 virus bearing the R5-tropic HIV-1 envelope JR-FL and similar results were obtained, indicating that the antiviral effects of Trim69 were envelope independent (**Figure 1G**).

In agreement with two previous reports [22, 23], Trim69 also potently modulated replication of the negative-strand RNA virus, VSV (**Figure 1H**). Next, we examined the ability of Trim69 to interfere with the replication of SARS-CoV2, a positive-strand RNA Coronavirus, in epithelial lung Calu-3 cells. Under these conditions, Calu3 expressing Trim69 were strongly protected from SARS-CoV2 infection (**Figure 1I**), overall indicating that Trim69 is able to interfere with a broad range of RNA viruses that extends from retroviruses to positive- and negative-strand RNA viruses (**extended data Figure 5** for non-normalized values of infection with the different viruses). Of note, Trim69 overexpression did not inhibit lentivirus infection in HEK293T cells, in agreement with one previous report [22]. Inhibition of VSV replication occurred in these cells, albeit to a lower magnitude with respect to what observed in the case of THP-1-PMA cells (**extended data Figure 6**), suggesting that the extent of the effects of Trim69 may be governed by a combination of both virus and cell type specific features.

### Trim69 inhibits HIV-1 reverse transcription and the early phases of VSV and SARS-CoV2 infection

To identify the step/s at which Trim69 interfered with virus replication, Trim69-overexpressing THP-1-PMA or Calu3 cells were challenged with the different viruses and the early phases specific to each one were analyzed (schematically resumed in **Figures 2A, 2B, 2C**). In the case of Lentiviruses, THP-1-PMA OE cells were challenged with R5-HIV-1 and virus entry into the cell was measured using the EURT assay (Entry/ Uncoating assay based on core-packaged RNA availability and Translation [31]), which is based on the direct translation of a Luciferase-bearing HIV-1 mini-genome incorporated into virion particles (**Figure 2A**). Under these conditions, Trim69 did not affect HIV-1 entry. On the contrary, Trim69 impaired the accumulation of all HIV-1 viral DNA intermediates tested (2.7 fold for MSSS to 5.3 for 2LTRs), indicating that this protein inhibits reverse transcription rapidly after the entry of viral capsids in target cells.

**Figure 2.**
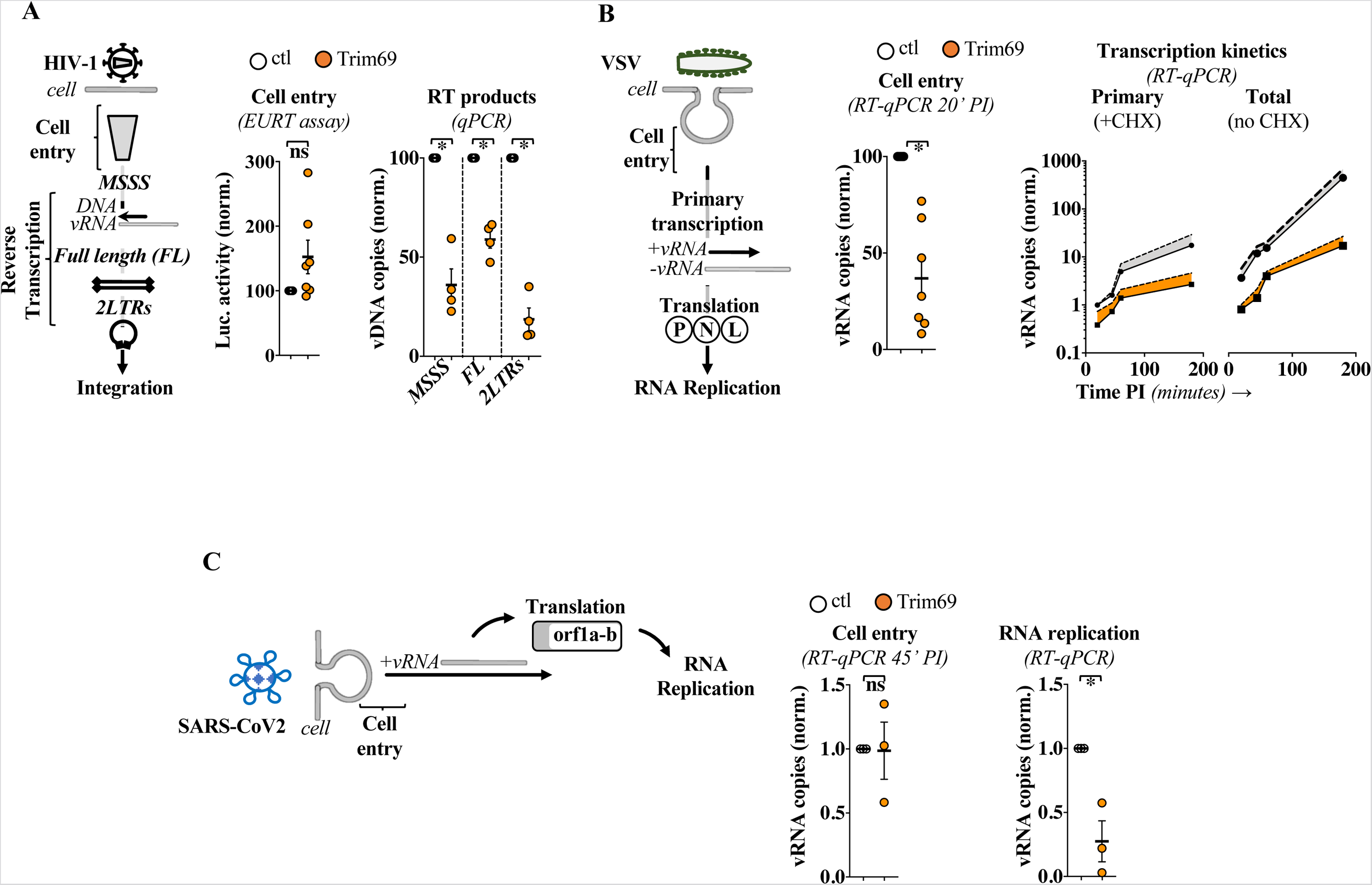
Trim69 inhibits the initial steps of infection of different viruses. A) As schematically presented, virus entry into target cells and accumulation of viral reverse transcription DNA intermediates were measured in THP-1-PMA cells overexpressing or not Trim69 upon challenge with an R5-tropic Env HIV-1. Cell entry was tested according to the EURT assay, an HIV-1-based cell entry assay based on the direct translation of an HIV genome mimic carrying the Firefly Luciferase. To ensure complete exposure of viral RNA, infections were carried out in the presence of 10 μM of PF74, prior to the quantification of the amount of produced luciferase 18 hours post viral challenge. B) VSV cell entry and RNA transcription steps were similarly tested in THP-1-PMA cells. Cell entry was quantified by measuring the levels of viral RNA that entered cells 20 min after viral challenge and trypsin treatment of target cells to remove unbound virus. Pioneer and whole RNA replication were discriminated by performing infections at an MOI of 1 in the presence or absence of 100 μg/mL of CHX. C) As in B for SARS-CoV2. The extent of infection was determined in Calu3 cells +/- Trim69 at 45 minutes post infection and after a trypsin treatment to remove non-internalized viruses. Viral RNA replication was then determined by RT-qPCR at six hours post infection to focus on early events. Graphs present Avg and SEM of 3 to 7 individual experiments. ns and *, non-significant and p<0.05 following a two-tailed Student t test between the indicated conditions.

VSV is a negative-strand RNA virus and, as such, it undergoes an obligate round of primary transcription after cell entry that is required for the translation of P, N and L proteins that in turn ignite viral RNA replication (for a review see [32] (**Figure 2B**). Given that translation from primary transcripts is required for viral RNA replication, the translation inhibitor cycloheximide (CHX) can be used to distinguish primary transcription from overall RNA replication. Contrarily to what was observed for HIV-1, TRIM69 exerted a measurable defect in VSV entry into the cell (2.9 fold) and this defect increased during primary transcription (6.5-fold) and overall RNA replication levels (34 fold). Thus in the case of VSV, Trim69 imparts successive cumulative antiviral effects.

SARS-CoV2 is a positive-strand RNA virus and as such its genome can be directly translated in viral proteins that ignite RNA replication (for a review see [33].

When Calu-3 cells overexpressing Trim69 were challenged with SARS-CoV2, no major defects were observed at entry, but a defect in RNA replication was clearly observable by six hours post infection (**Figure 2C**, 3.6 fold, in line with the replication defect observed).

Overall, these results indicate that Trim69 inhibits the early steps of viral replication of different viruses at slightly distinct steps: at a post entry step that affects the efficiency of reverse transcription and of viral RNA replication in the case of lentiviruses and of the SARS-CoV2 Coronavirus, and at both entry and post entry events in the case of VSV. In the latter, the efficiency of virus entry in cells expressing Trim69 exhibits a small but detectable defect, together with a more apparent defect in primary transcription.

### Trim69 is a novel regulator of stable microtubule dynamics

As a first step to decorticate the functions of Trim69, its intracellular distribution was determined by confocal microscopy. In agreement with a previous report [23], Trim69 adopts a filamentous distribution in the cell cytoplasm which is particularly marked in THP-1-PMA cells (**Figure 3A**). Exploratory analyses indicated that Trim69 exhibited little to no colocalization with a series of cellular markers in HEK293T cells (actin, ATG-3, ER, or CLIP170), with the exception of α-Tubulin (**extended data Figure 7**). Indeed, Trim69 colocalized strongly in all cell tested with stable microtubules (defined here as either acetylated or detyrosinated) which represent a subset of MTs (**Figure 3B**). Interestingly, Trim69-expressing cells exhibited higher levels of stable microtubules, suggesting that this protein could promote their formation.

**Figure 3.**
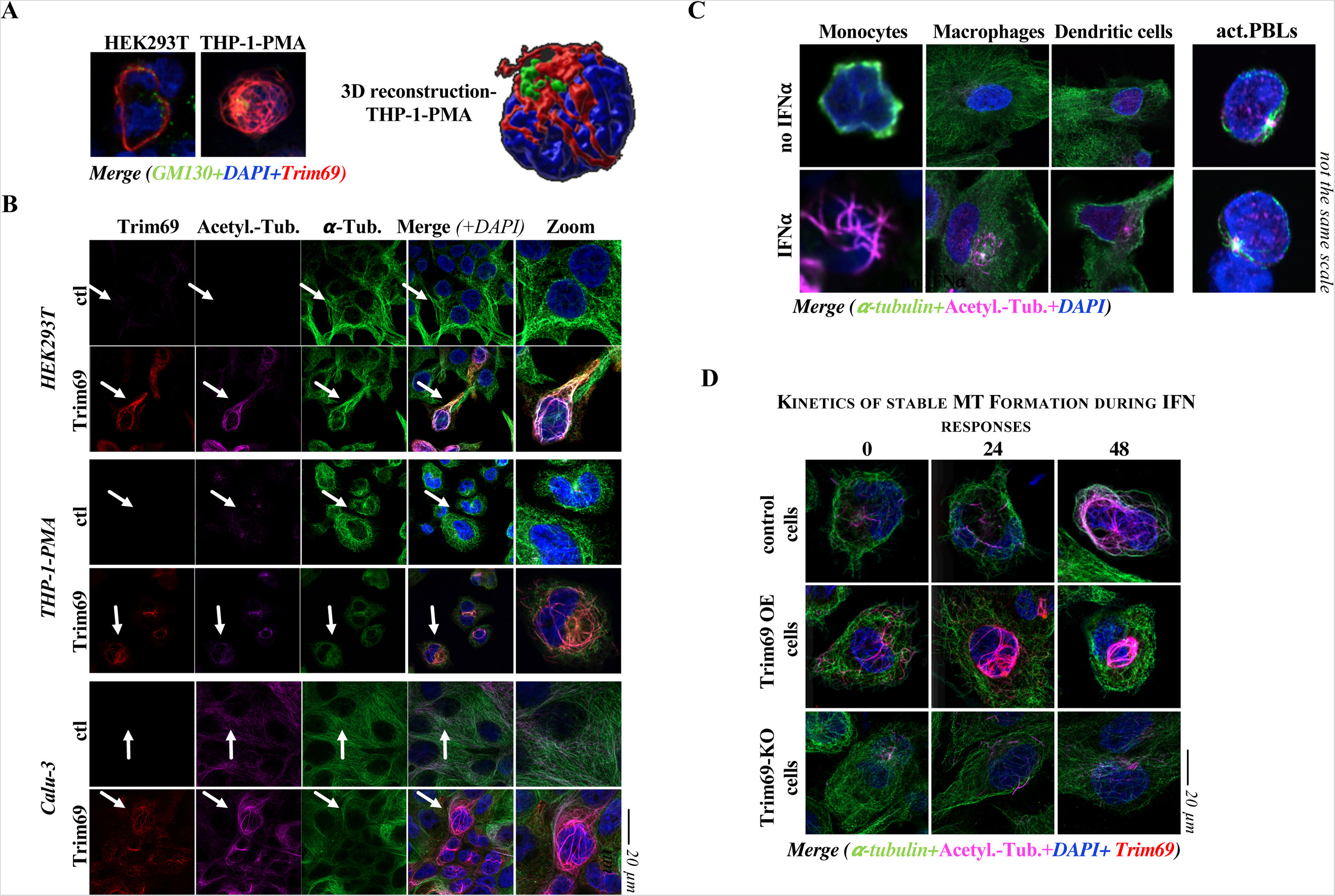
Trim69 is a novel and key mediator of IFN-I induced microtubule stabilization. A) Representative confocal microscopy pictures and 3D reconstruction of the intracellular distribution of Trim69 in HEK293T and THP-1-PMA cells. B) Distribution of Trim69 with antibodies recognizing endogenous α-tubulin or its acetylated form in the indicated cell types. Arrows indicates the cell zoomed on the right panels. C) Primary blood monocytes, macrophages, dendritic cells (DCs) as well as PH1/IL2 activated PBLs were stimulated or not with 1.000U/mL of IFNα for twenty-four and forty-eight hours prior to fixation and confocal microscopy analysis. Representative cells at 24 or 48 hours are presented here, while the quantification of stable microtubules, at all the time points is presented in the **extended data Figure 8**. Given the large differences in cell size, the pictures do not use the same scale. D) as in B in THP-1-PMA cells either control (*wild-type*) or in which Trim69 was either overexpressed or knocked out (OE and KO, respectively). Pictures present typical results obtained in >3 independent experiments in over 50 cells examined per condition. Separated channels for the images in panels C and D, in addition to the quantification of the amounts of stable MTs, are provided in the extended data Figures 8 and 9.

To put this observation in the context of IFN-I responses, we determined whether microtubule stabilization could be observed in response to IFN-I in different primary blood cells (**Figure 3C** and **extended data Figure 8** for separated channels and quantification of stable MTs on a per cell basis). Stimulation of primary blood monocytes as well as of monocyte-derived macrophages and dendritic cells (DCs) led to a pronounced upregulation of stable microtubules within 24-48 hours (monocytes>macrophages>DCs). In contrast, such accumulation was not observed in activated primary lymphocytes. Overall, these results indicate that IFN leads to a program of microtubule stabilization that may be particularly important for antiviral responses in cells of myeloid origins.

To determine the role that Trim69 may play in this program, we used THP-1, cells of myeloid origins more amenable to genetic manipulation. In control cells, stable MTs were present at low levels in the absence of IFN stimulation, but they increased over time following IFNα stimulation similarly to what was observed in primary macrophages (**Figure 3D** and **extended data Figure 9** for complete pictures and quantification of stable MTs on a per cell basis). As expected from our previous observations, expression of Trim69 (OE) increased the basal levels of stable microtubules accumulation and this was further stimulated by IFNα. However, the accumulation of stable microtubules was severely diminished in Trim69 KO cells stimulated with IFNα, indicating for the first time that the accumulation of stable microtubules is an integral part of the antiviral IFN response and that Trim69 plays an instrumental role in this program.

### Trim69 directly associates to microtubules

To determine whether Trim69 could physically associate to microtubules, a microtubule polymerization/sedimentation assay [34] was performed on THP-1 cell lysates expressing or not Trim69. In this assay, tubulin is induced into polymerization upon incubation with GTP and Taxol and stabilized microtubules are then purified by ultracentrifugation, along with associated cellular proteins (scheme of **Figure 4A**). Accordingly, a large fraction of microtubules sedimented in the pellet fraction upon Taxol stabilization and, when present, Trim69 was also present in this fraction, indicating that Trim69 was indeed physically associated to microtubules (**Figure 4A**, fraction P). To determine whether this association was direct, Trim69 was purified from bacteria as a fusion protein with the glutathione-S-transferase protein (GST). Commercially available and pure tubulin was then incubated in the presence of GTP and Taxol before binding to either GST or GST-Trim69 (**Figure 4B**). Under these conditions, Trim69 interacts with microtubules, indicating that Trim69 interacts directly with microtubules.

**Figure 4.**
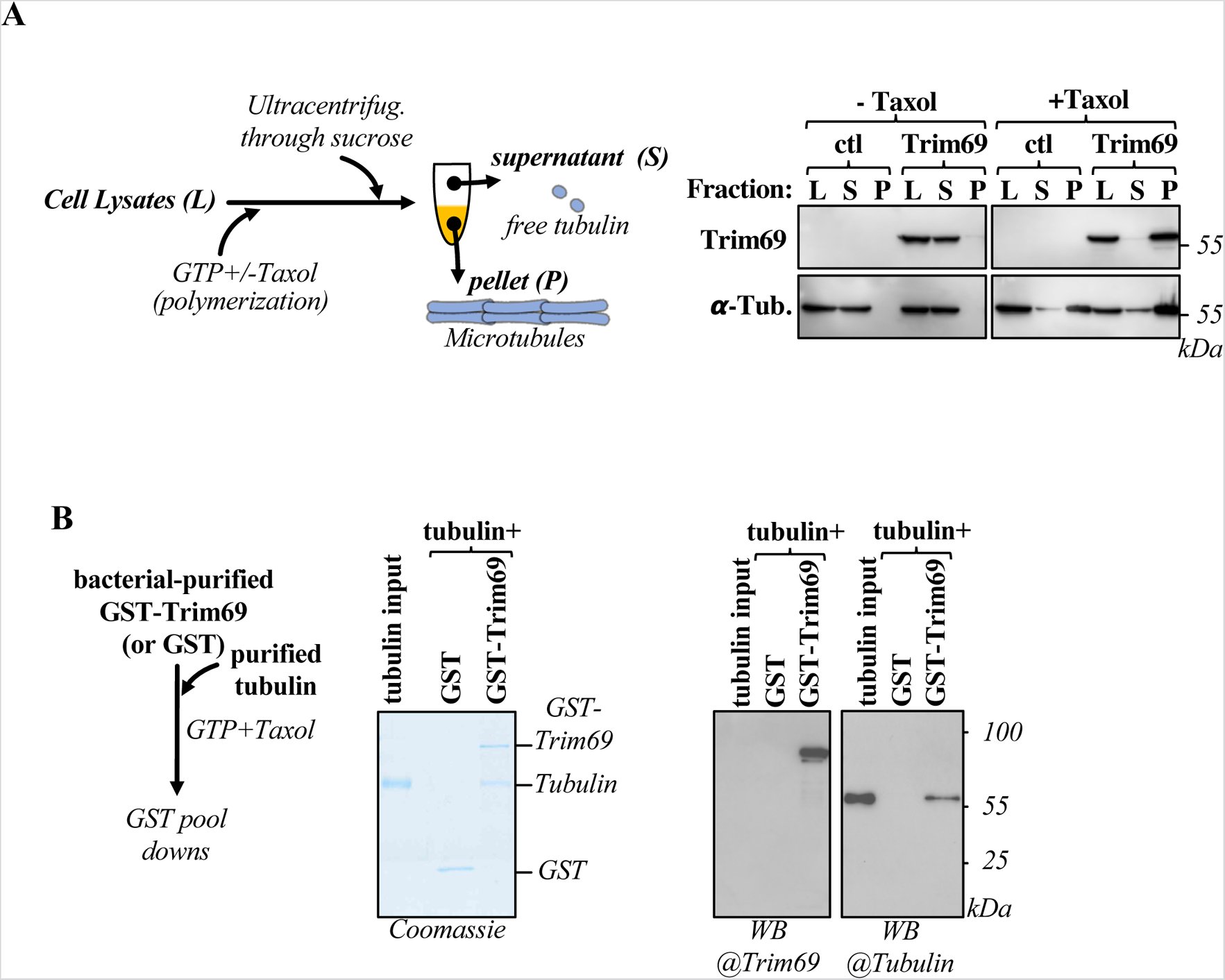
Trim69 associates directly to microtubules. A) Schematic representation of the microtubule sedimentation assay used. Briefly, the cytosolic fraction of THP-1-PMA cells expressing or not Trim69 was harvested and free cellular tubulin polymerization and stabilization was induced by incubation with GTP and Taxol. Microtubules and associated cellular proteins were then purified through a sucrose gradient prior to WB analysis. B) Direct binding between tubulin and Trim69, was assessed by using commercially available pure tubulin and GST-Trim69 purified from bacteria. The Coomassie and WB panels are representative of 3 independent experiments.

### Stable microtubule formation is key to the antiviral effects of Trim69

Five different isoforms issued from alternative splicing have been recently described for *trim69* (from the *wild-type* A to E) that contain extensive deletion in domains that are normally important for Trim family members functions. These isoforms were therefore expressed in THP-1-PMA cells prior to confocal microscopy analysis, or viral challenge with VSV (**Figure 5A** and **5B** and **extended data Figure 10** for WB analysis and separated IF channels). Under these conditions, isoforms B to E lost their ability to stimulate stable MTs and to protect target cells from viral challenge. Two additional mutations were then introduced in Trim69: mutations in two key cysteine residues that destabilize the RING domain (C61A/C64A), in addition to the deletion of the PRY-SPRY domain that in certain Trim members represents the domain of interaction with cellular partners (ΔP-SPRY, **Figure 5A** and **5B** and **extended data Figure 10**, as above). Similarly, to what was observed with the Trim69 isoforms, both mutants lost their ability to drive stable MT accumulation and both were unable to prevent viral infection. To gather further insights on the ability of the different mutants not only to stimulate stable microtubules, but also to co-localize with α-Tubulin, the Pearson’s coefficients between these two markers were determined (**Figure 5A**). Co-localization with α-Tubulin was lost for all mutants, suggesting that they have likely lost their ability to associate to microtubules in the first place.

**Figure 5.**
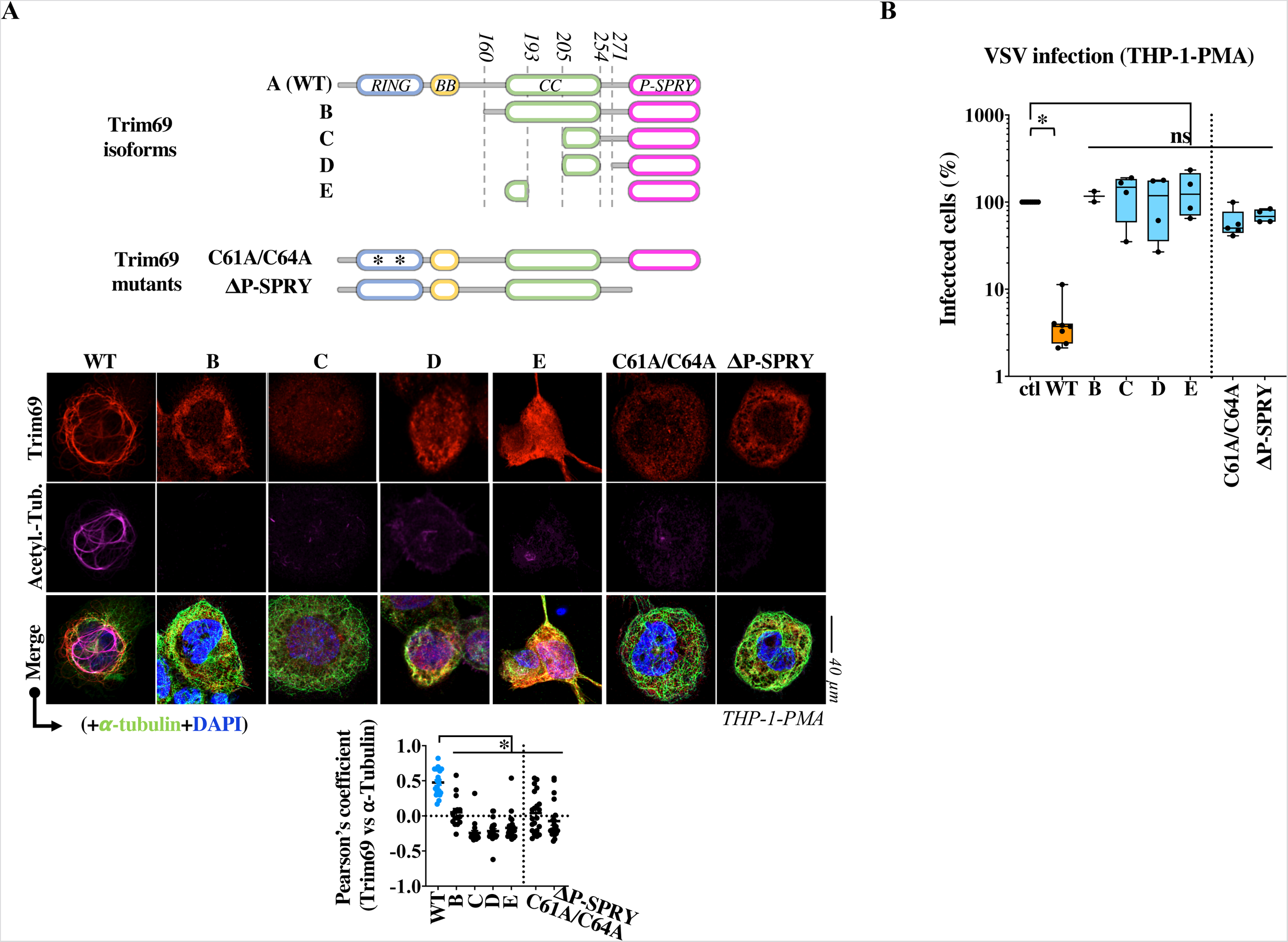
The ability of Trim69 to stimulate stable MTs is intimately linked to its antiviral abilities. A) Schematic presentation of Trim69 domains, isoforms and mutants used here. Mutants were evaluated for their ability to stimulate stable MTs formation in THP-PMA cells by confocal microscopy. The graph presents Pearson’s coefficients calculated between Trim69 mutants and α-Tubulin in 24 to 15 cells. *, indicates statistical significant differences between each mutant and WT, following an ordinary one-way Anova with Dunnett’s multiple comparison test. Representative WB panels of the different mutants in THP-1-PMA cells is provided in the **extended data Figure 10**. B) Cells were also challenged with an MOI of 0.1 of VSV, prior to flow cytometry analysis eighteen hours later. Pictures display representative patterns obtained and the box and whiskers presents data obtained from 2 to 4 independent experiments. Ns and *, respectively non significant and p=0.0005 following an ordinary one-way Anova with Dunnett’s multiple comparison test of the indicated samples over control.

Overall, these results indicate that the antiviral effects of Trim69 are intimately linked to its ability to drive the accumulation of stable MTs.

### Induction of stable MTs, as well as antiviral functions are conserved in a non-human divergent primate Trim69 ortholog

To increase the primate sequences available for phylogenetic analyses and to obtain a more complete evolutionary perspective of the selective pressure on Trim69 [20, 22], we *de novo* sequenced Trim69 from two additional monkey species, L’Hoest’s monkey (*Cercopithecus l’hoesti*) and Cotton-headed tamarin (*Saguinus oedipus*) (**Figure 7A**), and we determined the evolutionary history of Trim69 inferred from 26 primate species and 22 simian species (**Figure 7A, 7B** and **extended data Figure 11**). Additional species sequences and multiple methods to detect positive selection allowed us to confirm that Trim69 has been under adaptative evolution. The site-by-site analyses on the simian alignment allowed us to identify eight sites under positive selection distributed through the protein domains, with residues 158, 246 and 299 identified by several methods (**Figure 7A, 7B, 7C** and **extended data Figure 11**). Because the soTrim69 was one of the most divergent simian sequences from hTrim69 (88.6% identity; **Figure 7B**) and differed at 7/8 sites under positive selection, we cloned the soTrim69 gene to determine whether it exhibited distinct, or conserved, functionalities compared to its human counterpart. Stable THP-1-PMA cells expressing each of them were challenged with the indicated viruses prior to analysis by WB, confocal microscopy and flow cytometry to quantify the extent of infection (**Figure 7D**). Under these conditions, soTrim69 was capable of strong induction of stable microtubule formation and exhibited equivalent antiviral properties than hTrim69 upon challenge with VSV or distinct retroviruses. Overall, these results indicate that the main functional properties are maintained in the soTrim69 orthologue underscoring their importance in the antiviral properties of Trim69. Our data also indicate that retroviruses are unlikely to have driven positive selection in Trim69.

## DISCUSSION

In this work, we describe Trim69 as the first IFN-regulated protein that inhibits a diverse spectrum of viruses by promoting a global program of microtubule stabilization in the cell. By analyzing the response of several primary blood cell types to IFNα, we show that this program is a previously unrecognized facet of the innate defense system in cells of the myeloid lineage. Furthermore, using THP-1 cells that are more amenable to genetic manipulation, we show that Trim69 plays an instrumental role in this response. Trim69 interferes with model members of the *Retroviridae*, *Rhabdoviridae* and *Coronaviridae* families that collectively cover a large representation of replication modes amongst RNA viruses. The magnitude of inhibition depends on the combination between virus and cell type and in the cases of Lentiviruses, the antiviral effect is observed essentially in cells of the myeloid lineage. This is not unprecedented as other cellular factors inhibit HIV-1 in a cell type specific manner (for example SAMHD1, or APOBEC3A [35, 36]).

Viral inhibition occurs during the early phases of the different viruses’ life cycle, albeit with slight differences. In the case of HIV-1, inhibition occurs at reverse transcription, after entry of viral complexes into the cell and SARS-Cov2 seems to follow the same inhibitory path, with no measurable defects in virus entry, but an early defect in viral RNA replication. Instead, in the case of VSV a small but non negligible defect can be observed at the step of entry, which is followed by a first major detectable defect occurs during pioneer transcription, in line with a previous report [23].

Although it remains formally possible that Trim69 targets viral components, the very diversity of viruses examined here lends support to the hypothesis that Trim69 modifies the cellular environment in a manner that is preclusive to viral infection. The hypothesis we privilege is that Trim69 interferes with the movements of viral complexes along microtubules. Indeed, HIV-1 viral cores have been visualized as sliding along microtubules with inward rates of 1 μm/sec consistent with Dynein-dependent movement [37, 38] and adaptor proteins of this complex have been involved in this association (for example the Bicaudal D2 adaptor, BICD2, [39, 40]). Interestingly, interference with dynein-dependent movement has been associated to an early reverse transcription defect [41] which is consistent with the defects observed here for Trim69 [41]. In the case of Rhabdoviruses, the phosphoprotein P, which is part of the viral nucleoprotein complex along with the nucleocapsid protein (N) and the RNA polymerase (L), does interact with the dynein light chain 8 (LC8) [42–44] and Coronaviruses accumulation in the perinuclear region that evolves in double membrane vesicles (DMVs) is also influenced by dynein [45–47]. It is thus possible that dynein transport is inhibited in the presence of stable microtubules that are decorated with Trim69.

Several viruses have been described to induce stable MTs formation: Herpesviruses [48–50]; Influenza virus [51]; Hepatitis E virus [52]; Adenovirus [53, 54] as well as HIV-1 [55–57]. While in some studies the functional importance of stable MT formation remains unclear and according to our data could even represent a cellular response to viral infection, there are cases in which MT stabilization is associated to a pro-viral functionality. This seems to be the case for HIV-1 in which viral capsids that enter target cells induce prompt MT stabilization and mimic cellular cap-loading proteins to promote their loading onto MTs and their dynein-dependent transport towards the nucleus [55–57].

It is therefore counterintuitive that IFN may induce a program of MT stabilization via Trim69, as this particular pool of MTs exhibit higher stability and higher cargo trafficking propensity with respect to the dynamic pool of MTs [58, 59].

We believe that this dichotomy is however only apparent. First, while it is true that stable MTs can be preferentially used for cargo transport, the initial step of cargo loading is instead highly inefficient when it occurs on microtubules that are already detyrosinated (i.e. stable) [60]. As such, it is easy to envision that according to the timing at which microtubules become stabilized with respect to virus entry, MT stabilization can lead to either pro- or anti-viral outcomes.

Second, while stable MTs are often considered as a single homogeneous entity, the existence of several post-translational modifications on MTs, as well as the existence of numerous MT-bound factors, is likely to result in a far more complex functional heterogeneity of microtubules. In line with this contention, our study indicates that as part of an antiviral response, Trim69 starts a program of microtubule stabilization that lead to structures whose functionality is markedly antiviral.

Trim69 binds directly to MTs, but contrarily to other members of the Trim family we could not identify a C-terminal subgroup one signature (COS domain), a 60 amino acid stretch that mediates MT binding in certain Trim family members [61]. We then show that both its RING and PRY-SPRY domains are required for the antiviral activities of Trim69, as well as for microtubule stabilization. A most plausible model would therefore be that Trim69 associates to microtubules and that contacts relevant substrates via its PRY-SPRY domain leading to their degradation. However, mutations in the RING domain can also interfere with Trim homodimerization and the importance of the E3-ubiquitin ligase activity in the functions of Trim69 is debated [21, 23].

Finally, three studies including the present one have accumulated evidence of ongoing genetic conflict in primate Trim69 [20, 22], as well as of polymorphism in the human population. Using a very divergent simian Trim69, our results seem to exclude Lentiviruses that have invaded primates as main drivers of this selective pressure. However, Trim69 has been described to physically interact with the P protein of VSV and with the NS3 protein of Dengue virus [21–23], suggesting that viral antagonists of these or of other viral families that remain to be discovered may have exerted a genetic pressure on Trim69.

Overall, in a panorama filled with positive cofactors at the level of cytoskeleton, Trim69 represents for the moment the first antiviral factor that opposes viral infection by regulating microtubule dynamics. Given the fact that these structures influence several aspects of the cellular physiology, these findings may bear implications that extend beyond viral infection.

## METHODS

### Cells

Human embryonic kidney HEK293T cells (ATCC cat. CRL-3216), monocytic THP-1 cells (ATCC cat. Cat# TIB-202) and epithelial lung Calu-3 cells (ATCC cat. HTB-55) were respectively maintained in complete DMEM, RPMI-1640 or MEM media and 10% Fetal Calf Serum (FCS, Sigma cat. F7524). THP-1 cells media was also supplemented with 0.05 mM β-mercaptoethanol (Euromedex, cat. 4227-A) and 10 mM HEPES (Sigma, cat. H0887) and macrophage-like differentiation was induced upon a twenty-four hours treatment with 100ng/ml of phorbol 12-myristate 13-acetate (PMA) (Sigma cat. P1585). Calu-3 cells media was instead supplemented with a 1x final concentration of non-essential amino-acids (Sigma cat. M7145) and sodium pyruvate (Gibco cat. 11360-039).

Primary blood monocytes and lymphocytes were purified from blood leukopacks of healthy and anonymous donors, as described [62]. White leukocytes were first separated through a Ficoll gradient and then monocytes and lymphocytes enriched fractions were harvested at the interface and bottom of a Percoll gradient, respectively. Monocytes were further purified by negative depletion (monocyte isolation kit II, catalogue n° 130-091-153, Miltenyi; purity ≥ 90%) and when indicated, they were differentiated in either macrophages or dendritic cells (DCs) upon incubation for 4 to 6 days in complete RPMI-1640+ 10%FCS with either human Macrophage-Colony Stimulating Factor (M-CSF at 100 ng/mL; Eurobio, cat. 01-A0220-0050), or with Granulocyte-Macrophage-Colony Stimulating Factor and interleukin 4 [62] (GM-CSF and IL4, at 100 ng/mL each; Eurobio, cat. PCYT-221 and PCYT-211). PBLs were instead used without further purification. In this case, cell stimulation was carried out for twenty-four hours with 1 μg/mL of phytohemagglutinin (PHA, Sigma) plus 150U/mL of interleukin 2 (Eurobio, cat. PCYT-209). Unless otherwise specified, cells were incubated for twenty-four hours with 1,000 U/mL of human IFNα2 prior to use (Tebu Bio, cat. 11100–1). IFNβ was similarly used at 500 U/mL (R&D systems cat. 8499-IF-010).

### Antibodies

The following primary antibodies were used for WB, or confocal microscopy: mouse monoclonal antibodies; anti-α-Tubulin, anti-Flag (Sigma cat. T5168, cat. F3165, respectively) and anti-GM130 (BD biosciences cat. 556019 and cat. 610823); rabbit polyclonal antibodies: anti-GM130 (Abcam cat. Ab52649); anti-detyrosinated tubulin (Millipore cat. Ab3201) and anti-acetyl (K40)-α-tubulin (Abcam cat. ab179484) that were used interchangeably to label stable microtubules; goat anti-Flag polyclonal antibody (Biotechne cat. NB600-344). The following secondary antibodies were used for WB: anti-mouse, anti-rabbit IgG-Peroxidase conjugated (Sigma, cat. A9044 and cat. AP188P); while the following ones were used for confocal microscopy: donkey anti-rabbit IgG–Alexa Fluor 594 conjugate and donkey anti-mouse IgG–Alexa Fluor 488 conjugate (Life Technologies, cat. A-21207 and cat. A-21202) donkey anti-rabbit IgG–Alexa Fluor 555 conjugate (Life Technologies, cat. A32794); donkey anti-rabbit IgG– Alexa Fluor 647 conjugate (Life Technologies, cat. A-21447); donkey anti-goat IgG–Alexa Fluor 546 conjugate (Life Technologies, cat. A-11035).

### Plasmid DNAs, retroviral vectors and viruses

Full length Trim69 (gene ID: 140691, isoform A) was obtained by gene synthesis and codon optimization (Genewiz) and it was cloned with an N-terminal Flag-tag in pRetroX-tight, a murine leukemia virus (MLV)-based retroviral vector that can be used for the generation of stable cells in which the expression of the gene of interest is under the control of doxycycline (dox. Ozyme, cat. 631311). Trim69 mutants were generated by standard molecular biology techniques. DNAs coding for several cellular markers were obtained through Addgene and in particular: mCherry-Actin-C-18, mEmerald-CLIP170-N-18 and mCherry-ATG3-C-18 were gifts from Michael Davidson (Addgene cat. 54967, 54044 and 54993) [63]; mCh-alpha-tubulin was a gift from Gia Voeltz (Addgene cat. 49149), while pcDNA-D1ER was a gift from Amy Palmer & Roger Tsien (Addgene cat. 36325). For presentation simplicity, the cellular markers cited above and Trim69 were pseudocolored in green and red, respectively, in the qualitative colocalization analysis presented in the **extended data Figure 7**.

Single round of infection competent HIV-1, HIV-2 and SIV_MAC_ lentiviruses have been described before [64]. Briefly, viruses were generated upon transient transfection of HEK293T cells with DNAs coding three components: the structural Gag-Pro-Pol protein of the virus of interest, the envelope protein (in our case either the G protein of the Vesicular Stomatitis Virus, VSVg, or the HIV-1 R5 Envelope, JR-FL) and a mini-viral genome containing a reporter gene expression cassette (CMV-GFP and for the HIV-1 screening the Firefly luciferase (obtained from Didier Negre, CIRI-Lyon, France). An MLV- or an HIV-1-based vector system were used for the generation of stable cell lines overexpressing Trim69 under the control of doxycycline (OE), or to generate *trim69* knockout cells (KO) by CRISPR/Cas9-mediated gene targeting, respectively.

Replicative VSV virus (Indiana serotype) bearing a GFP reporter transcription unit inserted between M and G have been described before [65]. Replicative SARS-CoV2 virus (Wuhan strain, [66]) bearing the mNeonGreen reporter inserted into the ORF7 was obtained from Pei-Yong Shi from the University of Texas Medical Branch, Galveston, TX, USA.

#### De novo sequencing of cotton-headed tamarin (Saguinus oedipus) and L’Hoest monkey (Cercopithecus l’hoesti) Trim69

First, simian peripheral blood mononuclear cells (PBMCs) were isolated using Histopaque 1077 from leftover blood samples (approximately 1 ml, from blood drawn for health purposes) from a L’Hoest monkey (Old World monkey *Cercopithecus l’hoesti*) hosted at the Zoo de Lyon, France (Guillaume Douay). Second, B92a cells (a gift from Branka Horvat, CIRI Lyon) from cotton-headed tamarin (New World monkey *Saguinus oedipus* were maintained in Dulbecco’s modified Eagle’s medium (DMEM) supplemented with 10% fetal calf serum. Total RNA was extracted from cells following the manufacturer’s instructions (Macherey-Nagel NucleoSpin RNA cat. 740956). Reverse transcription was performed using SuperScript III reverse transcriptase (Thermo Fisher cat. 18080) with random hexamers and oligo(dT). Single-round PCR was performed using Q5 high-fidelity DNA polymerase (NEB; M0491) following the manufacturer’s instructions, with primers targeting the UTRs: T69-F1: 5’-TCATGCTCTGAGYYCATTCC, and T69-R1: 5’-TCAATACCTCTTTAATAWCACTCTG. The sequences of the TRIM69 gene from cotton-headed monkey (soTrim69) and from L’Hoest monkey (cerLhoTrim69) are available at Genbank under accession numbers ON745601 and ON745600, respectively. The coding region of soTRIM69 was subsequently cloned in the pRetroX-tight using standard molecular biology techniques.

### Viruses production and infections

Single round of infection competent HIV-1, SIV_MAC_ or HIV-2 viruses were produced by calcium phosphate DNA transfection of HEK293T cells (Gag-Pro-Pol packaging construct, mini-viral genome and pantropic VSVg envelope: ratio of 8:8:4, for a total of 20 μg per 10cm dish). In the case of HIV-1 viruses bearing the HIV-1 JR-FL envelope, the ratio used was: 8:8:2: plus 0.5 of HIV-1 Rev to stimulate Env production). Supernatants were collected at forty-eight hours post transfection, syringe-filtered to remove cellular debris and virions were purified through a 25(w/v) sucrose cushion for 1h15min at 28.000 rpm. The number of infectious viral particles was determined by infecting HeLa cells with virus dilutions and by quantifying the number of green fluorescent protein (GFP)-positive cells obtained two-three days after by flow cytometry. The infectious titer equivalent of non-GFP coding viruses was determined against standards of known infectivity by exo-RT or p24 ELISA [64].

VSV and SARS-CoV2 viral stocks were produced by infecting HEK293T cells (10cm plates) or VeroE6 cells (T125 flasks) and by harvesting cell supernatants obtained 18 to 36 hours later. Infectious titers were measured by limiting dilutions infections followed by flow cytometry analysis 18 hours post infection. Infections were carried out with multiplicities of infection (MOIs) comprised between 0,1 and 5 depending on the viruses, as specified in the Figure legends. The percentage of infected cells was determined at different times post infection, depending on the virus: retroviruses (two days); VSV (eighteen hours); SARS-Cov2 (two days). In all cases, infected cells were fixed and analyzed by flow cytometry on a FACSCantoII (Becton Dickinson, USA).

### Generation of stable cell lines

Doxycycline-inducible stable cells overexpressing Trim69 were generated using the pRetroX-Tight system (Clontech), a murine leukemia virus (MLV) retroviral-based gene transduction system, as described earlier [67]. Briefly, MLV retroviral vectors were produced by calcium phosphate DNA transfection of HEK293T cells with plasmids coding MLV Gag-Pro-Pol, the VSVg envelope and two pRetro-X based mini-viral genomes, the first bearing Trim69 under the control of the dox-inducible promoter and the second coding the transcriptional transactivator rtTA (ratio 8:4:4:4 for a total of 20 μg per 10cm dish). Virion particles in the cell supernatant were directly used for cell transduction followed by selection of cell pools thanks to the Puromycin (Sigma, cat. P8833) and G418 (Sigma, cat. G8168) resistance genes carried by the two constructs. Stable cell lines were freshly generated every two-weeks maximum.

CRISPR/Cas9 *trim69* KO pools of stable cells were generated by two successive rounds of gene transduction with Cas9 and CRISPR-bearing vectors (gift of Feng Zhang, obtained from Addgene cat. 52963 and cat. 52962). In this case, selection of pool cells was carried out thanks to the Blasticidin (Invivogen cat. ant-bl-1) and Puromycin (Sigma cat. P8833) resistance genes carried by the two vectors. Two guides were used as described in [22]. Two guides were cloned into the BsmBI site of LentiGuide-Puro by annealing/ligation of two overlapping oligonucleotides (up and down, respectively): Guide 1, nt 276-295 of coding sequence, CACCGCAACCCTGTACTGGACAAGT and AAACACTTGTCCAGTACAGGGTTGC; Guide 2, nt 311-329 of coding sequence, CACCGAAGAAGTTACCCTTACTCAA and AAACTTGAGTAAGGGTAACTTCTTC. Effective gene editing was determined by PCR on genomic DNA using primers CACTTTCAAAGGAGAGATTATGTGC and GAGCAGTCTGGGCTTTCTAAT, cloning of the corresponding PCR fragment and sequencing of 10 individual clones.

### Library details and screening procedure

The shRNA library originated from the Broad institute Genetic Perturbation platform and individual shRNA clones were purchased from Sigma in an pLKO or pLKO_TRC005 context (3 shRNA per genes for a total of 419 genes, list provided in the supplementary file). shRNA coding HIV-1 vectors were prepared in 96 well plates by ectopic transfection of HEK293T cells by calcium phosphate DNA transfection of 80ng of Gag-Pro-Pol, 80 ng of the pool of the 3 shRNA coding constructs, plus 40 ng VSVg. Cycling THP-1 cells were transduced with three rounds of HIV-1 coding LVs preparations (daily), prior to puromycin selection (at 2.5 μg/mL). Cells were then counted, split in two and then differentiated for 24 hours with PMA prior to an additional 24 hours incubation period with or without 1.000 U/mL of IFNα. Cells were then challenged with an MOI equivalent of 0.5 of VSVg-pseudotyped HIV-1 virus coding the Firefly Luciferase reporter, prior to cell lysis and Luc analysis forty-eight hours later (Promega cat. E4530, according to the manufacturers’ instructions). An IFN defect was calculated for each gene as a ratio of the Luc. activities measured in the no IFNα/+IFNα conditions, normalized to the average value obtained for all genes together. Retained changes were below/above 2.5, with a statistically significant Student t test value over control. Retrieved hits that led to either a relieved or increased IFN defect were then recategorized as pro- or anti-viral proteins, according to their behavior in the no IFNα condition, as the IFN defect ratio is also influenced by this parameter. CD40 and TNPO3 were added as sentinel samples for routine control of silencing efficiency and effect under library screening conditions.

A secondary screen was performed to exclude primary hits whose silencing was likely to alter HIV-1 infection indirectly, by modulating IFN signaling. To this end, shRNA stable cells were differentiated with PMA and stimulated with 1.000 U/mL of IFNα. IP10 secretion in the supernatant was measured by ELISA, according to the manufacturer’s instructions (R&D cat. DY266).

### Phylogenetic and positive selection screen using DGINN

The Consensus CoDing Sequences (CCDS) were downloaded from the CCDS database (https://www.ncbi.nlm.nih.gov/CCDS/) using the CCDSQuery script provided with DGINN, with the exception of LILRA3 for which the coding sequence was manually retrieved from the NCBI databases (human sequence NM_006865) as no CCDS existed. In cases where multiple CCDS were referenced for one gene, the longest one was kept. We used the DGINN pipeline ([24]; available at: http://bioweb.me/DGINN-github) for the analyses in a two-step fashion. For the primate homologous sequence retrieval and phylogenetic analyses, we used the NCBI nr database limited to primate species, with otherwise default parameters (blastn e-value 10-4, identity 70% and coverage 50%, at least eight species for separation of orthologous groups). Alignments and phylogenetic trees produced during this first step were then used for positive selection analyses, according to different packages: BUSTED and MEME from the Hyphy package [68]; Codeml from PAML [69] and Bio++ [70], all running codon substitution M1, M7 and M2, M8 models that do not and do allow codons to evolve under positive selection, respectively. BUSTED and MEME from the Hyphy package [68] to look for gene-wide and site-specific episodic positive selection. Genes were considered under positive selection for a BUSTED p-value < 0.05 and sites for a MEME p-value < 0.05. Codeml from PAML [69] and Bio++ [70] were used to run codon substitution models M1, M2, M7 and M8. M1 and M7 are neutral models not allowing for positive selection and M2 and M8 are their pendant with a class allowing codons to evolve under positive selection. P-values derived from the likelihood ratio tests between the two models (M1 vs M2, M7 vs M8) were used to determine which model is a better fit for the data. Genes were considered under positive selection for p < 0.05 and sites for posterior probabilities (PP) > 0.95 (in the Bayes empirical bayes BEB test for Codeml M2 and M8, and in the PP test for bio++ M2NS and M8NS).

#### Phylogenetic and positive selection analyses of primate and simian TRIM69

The codon alignment of primate TRIM69 from the DGINN screen was retrieved, and new sequences from NCBI databases and from de novo sequencing were added. Codon alignments of primate TRIM69 sequences and simian-only TRIM69 sequences were performed using MUSCLE [71] and manually edited, resulting in 26 and 22 included species, respectively. Phylogenetic analyses were performed with PhyML with a GTR+I+G model and 1,000 bootstrap replicates [72]. Positive selection analyses were performed in DGINN using the new codon alignments, as previously described. Additionally, FUBAR and MEME were run in Datamonkey [73] and sites with PP>0.9 and p<0.1 were considered under positive selection, respectively.

### Analyses of the virus life cycle steps affected by Trim69

#### HIV-1 entry (<u>E</u>ntry/ <u>U</u>ncoating assay based on core-packaged <u>R</u>NA availability and <u>T</u>ranslation system, EURT)

This assay is based on the direct translation of a Luciferase-bearing HIV-1 genome mimic that can be directly translated, yielding a precise measure of virus entry into the cell cytoplasm [31]. Single round of infection R5 Env HIV-1 virus incorporating this reporter (EU-repRNA) were produced by transient transfection of HEK293T cells, as described above. Infections were carried out overnight in THP-1-PMA cells expressing or not Trim69, with a MOI-equivalent of 2, prior to cell lysis and Luciferase activity quantification (Promega, cat. E2620, according to the manufacturers’ instructions). To increase the levels of capsid opening, the capsid destabilizing compound PF74 was added for the assay at 10 μg/mL (Sigma, cat. SML0835-5).

#### HIV-1 reverse transcription

The quantification of revere transcription vDNA intermediates was performed as described earlier [13]. Briefly, cells were infected with DNase-treated R5-HIV-1 virus preparations at an MOI of 2 to 4 and lysed 24h post infection, and extracted DNA was used to amplify the different vDNA forms with specific primers (from 5’ to 3’): minus-strand strong stop (MSSS), TGGGAGCTCTCTGCTAACT and ACCAGAGTCACACAACAGACG; full-length (FL-GFP), GAACGGCATCAAGGTGAACT and TGCTCAGGTAGTGGTTGTCG; HIV-1 2LTR, TCGTTGGGAGTGAATTAGCC and CCCACTGCTTAAGCCTCAAT. Cellular HPRT or 18S RNA were used for sample normalization: TGACCTTGATTTATTTTGCATACC and CGAGCAAGACGTTCAGTCCT, or GTGGAGCGATTTGTCTGGTT and CGCTGAGCCAGTCAGTGTAG, respectively.

#### VSV entry and RNA replication

VSV entry was measured by quantifying the levels of viral RNA in cells 20 min after infection by RT-qPCR. Prior to cell lysis, cells were extensively washed and treated for 10 minutes at 37°C with trypsin to remove non-internalized virions. Pioneer and overall RNA replication levels were measured by performing infections at a MOI of 0,1 to 0,5 for different times points in the presence or absence of 100 μg/mL of chycloheximide (CHX, Sigma, cat. 4859). This compound blocks the translation of viral proteins from positive strand viral RNAs, preventing the accumulation of proteins required for the full replication cycle of RNA. It thus allows the distinction between pioneer transcription and full RNA replication. Primers amplified the GFP gene inserted into the viral genome: GAACGGCATCAAGGTGAACT and TGCTCAGGTAGTGGTTGTCG.

#### SARS-CoV-2 entry and RNA replication

SARS-CoV-2 entry was measured as described above by measuring the amount of viral RNA inside cells 45 min after infection. Variations in the amount of viral RNA were then assessed at an early time point after infection by RT-qPCR (6 hours), as pilot experiments indicated defects in viral RNA replication in the presence of Trim69 already at early time points. Primers amplified the Neongreen gene inserted into Orf7 were: GAGCTGCATATCTTCGGATCCATCAACG and CAGGTCTCCCTTGGTACTCTTCAGGTTC.

#### RNA extractions and RT-qPCR

Total cellular RNA was extracted according to the manufacturer’s instructions (Macherey-Nagel NucleoSpin RNA cat. 740956) and reverse transcription was performed using random hexamers and oligo(dT) with the SuperScript III reverse transcriptase (Thermo Fisher cat. 18080) also following the manufacturers’ instructions. qPCRs were performed on a StepOne Plus real-time PCR system (Applied Biosystems) using the FastStart universal SYBR green master mix (Roche Diagnostics, cat. 4913914001). For the quantification of the levels of Trim69 mRNA levels, the PCR primers used were as published (Wang et al, 2018) (TCTGTGGGGCAGTCTAAGGA and CCATGGACACATGTTGCTGC) and HPRT1 (see above) was used to normalize samples.

### Microtubule binding assays

#### Microtubule sedimentation in THP-1-PMA cell lysates

The protocol was adapted from [34]. Briefly, THP-1-PMA cells in which expression of Trim69 had been induced for forty-eight hours with 0.5μg/ml dox were lysed in PEM buffer (80 mM PIPES pH 6.8, 1mM EGTA, 1mM MgCl_2_, 0.5mM DTT,150mM NaCl,1% IGEPAL) supplemented with protease inhibitors (Sigma, cat. 4693159001), at 4°C for 1 hour. Cell lysates were then depleted of insoluble material by four consecutive rounds of centrifugation (610g for 10 min; 10.000g for 10 min; 21.000g for 20 min and last at 100.000g for 1 hour all at 4°C). The resulting cell lysate (L) was then supplemented with 2 mM GTP +/- 40 μM Taxol (Life technologies, cat. R0461 and P3456) and incubated at 37°C for 30 min to induce the polymerization of free tubulin into microtubules. Samples were then layered over a 15% (w/v) sucrose cushion and centrifuged at 30.000 g for 30 min at 30°C to sediment polymerized microtubules (P) and associated proteins from unpolymerized tubulin (S). All samples were then equalized in sample buffer prior to immunoblot analysis.

#### *In vitro* binding experiments

Flag-Trim69 was cloned downstream of a gluthathione-S-transferase (GST) in the pTKPL vector (an in-house derivative of the vector pGEXTK, Promega) and purified from bacterial lysates on glutathione agarose beads (Sigma, cat. 17-0756-01). Equal amounts of GST or GST-Trim69 proteins were then bound to 10 μg of pure porcine brain tubulin (purchased from Euromedex, cat. CS-T240-A) in a total volume of 40 μl of PEM buffer supplemented with 40 μM of Taxol and 1mM GTP. Samples were first incubated for 45 min at 37°C and then incubated for 1 hour at 4°C to induce microtubule polymerization first and Trim69 binding, respectively. Beads were washed three times in the same buffer prior to analysis.

### Confocal microscopy

Cells were plated on 0.01% poly-L-lysine-coated coverslips (Sigma, cat. P4832) and analyzed 24 hours after ectopic DNA transfection (unless otherwise specified, Lipofectamine 3000 cat. L3000008, ThermoFisher, according to the manufacturer’s instructions). Cells were washed three times with PBS 1x, fixed with 4% paraformaldehyde (Euromedex, cat. 15713), quenched with 50 mM NH_4_Cl (Sigma cat. A4514) and permeabilized with PBS–0.5% Triton X-100 (Sigma, cat. X100) (the timing of these steps was 10, 10 and 5 minutes each). After a blocking step in PBS–5% milk, cells were incubated with primary antibodies for 1 hour at room temperature (dilution 1:100), washed and then incubated with fluorescent secondary antibodies (dilution 1:100). A 4’-5-diamidina-2-phenylindole (DAPI)-containing mounting medium was used (ThermoFisher, cat. 62248). Images were acquired using a spectral Zeiss LSM800 confocal microscope or Confocal Zeiss LSM980 - AiryScan and analyzed with Fiji software (version 2.0.0). Colocalisations were quantified using the Pearson’s overlap coefficient, and intensities of stable tubulin were quantified by measuring the pixels Integrated Density function weighted for the area in each cell (Fiji software).

### Softwares

Confocal microscopy: Fiji software (version 2.0.0), Zen (version 2.3, Zeiss) and Imaris 9.2.0 software (Oxford Instruments Group). WB: Image Lab Touch Software (version 2.0.0.27, Chemidoc Imaging System from Bio-Rad). Flow cytometry: FlowJo (version X, BD). Protein structure modeling: RaptorX (http://raptorx.uchicago.edu/) [74].

Statistics and graphs: Graphpad Prism8 (8.4.3, Graphpad software, LLC) and Excel (16.16.3, Microsoft).

### Statistical analyses

Statistical analyses were calculated with the Graphpad Prism8, or Excel softwares: Student t tests (unpaired, two-tailed), one-way Anova tests with either Tukey’s or Dunnett’s multiple comparisons, as indicated.

### Ethics statement

Primary blood cells were obtained from the blood of healthy donors (EFS-Lyon) in the form of discarded “leukopacks” obtained anonymously so that gender, race, and age of donors are unknown to the investigator and inclusion of women, minorities or children cannot be determined. The work was carried out under the authorization n° 19-606 from the Ethical Commission of the INSERM (CEEI) and n° DC-2019-3774 CODECOH from the French Ministry of Research.

### Data availability

Source data is provided with this paper. There is no restriction on data availability. Further information and requests for resources and reagents should be directed to the Lead Contact, Andrea Cimarelli. All reagents generated in this study are available upon request from the Lead Contact with a completed Materials Transfer Agreement.

## Acknowledgements

We thank Laurent Guéguen at the LBBE, Lyon, for fruitful discussions and scientific insights on the phylogenetic and positive selection analyses. We acknowledge the contribution of the microscopy (LYMIC-PLATIM) platform of SFR BioSciences Gerland Lyon Sud (UMS3444/US8). We wish to thank Didier Négre and Philippe Mangeot at the CIRI and Pei-Yong Shi from the University of Texas Medical Branch, Galveston, TX, USA for sharing of material, in addition to the authors that provided material through Addgene. We thank Branka Horvat (CIRI, Lyon) for sharing B95a cells, as well as Guillaume Douay and the Zoo de Lyon for their collaboration and leftover blood samples from hosted monkeys. We thank all the contributors of publicly-available genomic and genetic sequences, as well as of phylogenetic programs. YS is the recipients of a PhD fellowship from the Chinese Scholarship Council (CSC); AK and CdS have been the recipients of a post-doctoral fellowship from the ANRS|Maladies infectieuses émergentes (ANRS|MIE). This work has been supported by grants from the ANRS (AO-2014-1 and AO-2021-1) to AC, from the French Research Agency on HIV and Emerging Infectious Diseases ANRS/MIE (no. ECTZ19143 and ECTZ118944) and the ANR LABEX ECOFECT (ANR-11-LABX-0048 of the Université de Lyon, within the program Investissements d’Avenir [ANR-11-IDEX-0007]) to LE. AC and LE are researchers of the Centre National de la Recherche Scientifique (CNRS). The funders had no role in study design, data collection and analysis, decision to publish, or preparation of the manuscript.

## Author Contributions

Y.S. designed and performed most of the experiments. XNN, carried out experiments with CRISPR/Cas9 KO cells and HIV infection, while CdS performed the initial genetic screen and AK the initial characterization of Trim69‡. L.P. and L.E. performed the evolutionary analyses and provided monkey samples. A.C. supervised the research and wrote the manuscript. All authors commented on the manuscript.

## EXTENDED DATA

**Extended data Figure 1. Zoom on the infectivity results and recategorization of functionally retrieved hits.** A) Primary IFNα screen hits were defined as genes that when silenced modulated the normalized IFN defect above or below the chosen threshold value (2.5 fold), in a statistically significant manner when compared to the shRNA control sample, following an unpaired Student t test. Given that the IFN defect ratio is also influenced by the extent of infection in untreated cells (no IFN condition), which is itself dependent on the relative gene silencing efficiency and differential gene expression between basal and IFN conditions, the IFN defect cannot be used directly to categorize primary hits as pro- or anti-viral factors. To do so, the infectivities obtained upon gene silencing in the absence of IFN was used to categories primary hits as pro- or anti-viral modulators of infection (B). Genes that upon silencing relieved infection in the absence of IFN in a statistically significant manner were thus recategorized as antiviral (AIM2, TOP1, IGF2BP2 and ABTB2), while on the contrary, genes that upon silencing decreased the extent of infection in unstimulated cells were classified as pro-viral (TNPO3 and AKAP2). Remaining genes did not exhibit statistically significant changes among the two conditions and were not recategorized. The final list of retrieved hits and their respective categories are presented in **Figure 1D**.

**Extended data Figure 2. Complementary DGINN analyses of retrieved genes.** A) The graph presents the percent of residues under positive selection on the indicated hits, following DGINN analysis. B) Example of DGINN output and positive selection analyses in the indicated genes. Height of the bars indicates the number of methods that detected positive selection at the indicated position. C) DGINN automatically-detected paralogues and gene recombination events.

**Extended data Figure 3. Homologies between Trim69 and other members of the Trim family.** A) Trim69 amino acid sequences were used as query for a protein Basic Local Alignment Search Tool (BLAST) analysis. Blue hit map depicting percentage identities between the indicated Trim members using either the entire protein, or its individual domains. B) Overlay between the modeled 3D structures of the PRY-SPRY domains of human Trim5α and Trim69 (in red and grey respectively, RaptorX). V1 to V3 indicate variable loops known to be important for Trim5α target recognition.

**Extended data Figure 4. Trim69 cell expression pattern and effects of stable overexpression/knock out of Trim69 on cell division.** A) The indicated cells were stimulated with 1.000 U/mL of IFNα or with 500 U/mL of IFNβ for twenty-four hours, prior to RNA extraction. The levels of expression of Trim69 were determined by RT-qPCR (Avg and SEM of > 3 independent experiments/donors). Primary macrophages were differentiated from blood monocytes with M-CSF, while PBLs were activated for 24 hours with PHA/IL2. B) THP-1 cells either expressing Trim69 under the control of doxycycline (OE), or knocked out for Trim69 (KO) were induced with 1 μg/mL of dox. and then plated at equal density. Cell division was measured at the indicated time. The graph presents values obtained in three independent experiments.

**Extended data Figure 5. Trim69 expression in stable cells and non-normalized values of infection.** A) Representative results of Trim69 expression in dox.-inducible cells, as assessed by WB. B) The graph presents the non-normalized values of cells infected with the indicated viruses in Fig 1.

**Extended data Figure 6. Trim69 does not inhibit lentiviral infection in HEK293T cells.** Trim69 was ectopically expressed in HEK293T cells upon calcium phosphate DNA transfection and cells were then challenged with the indicated virus at an MOI comprised between 0.1 and 0.5 prior to flow cytometry analysis 18 hours (VSV) or 3 days later (Lentiviruses). The graph present Avg and SEM obtained in 3 to 9 independent experiments. * indicates p<0.05 following a two-tailed Student t test between the indicated conditions.

**Extended data Figure 7. Qualitative confocal microscopy assessment of the colocalization of Trim69 with known cellular markers.** HEK293T cells were ectopically transfected with DNAs coding TRIM69 along with the indicated fluorescent cellular markers (all colored or pseudo-colored in green for simplicity), prior to confocal microscopy analyses. The right panels present typical Pearson’s correlation coefficients between Trim69 and the indicated marker, as indicated.

**Extended data Figure 8. Complete confocal microscopy panels of the analysis presented in Figure 3C and stable MTs quantification.** Separated channel panels, not presented in the main figure for lack of space, are presented here. The graph presents the quantification of the amount of stable tubulin on a per cell basis (Integrated Density weighted for area, IntDen/area) calculated on 30 cells per condition/time point. * and ns, indicates statistically significant, or non-significant, following an ordinary one-way Anova with Tukey’s multiple comparisons test, as indicated.

**Extended data Figure 9. Complete confocal microscopy panels of the analysis presented in Figure 3D and stable MTs quantification.** Separated channel panels, not presented in the main figure for lack of space, are presented here. This figure also displays results obtained with control KO THP-1 cells. The graph presents the quantification of the amount of stable tubulin on a per cell basis (Integrated Density weighted for area, IntDen/area) calculated on 27 to 32 cells per condition. * and ns, indicates statistically significant, or non-significant, following an ordinary one-way Anova with Dunnett’s multiple comparisons test.

**Extended data Figure 10. Complete confocal microscopy panels of the analysis presented in Figure 5A**. Separated channel panels, not presented in the main figure for lack of space, are presented here along with representative WB analysis of the different mutants.

**Extended data Figure 11. Trim69 evolutionary analyses.** A) Phylogenetic tree of primate Trim69 corresponding to **Figure 6A**, with the NCBI reference of the sequences and the species nomenclature (three first letters from genus and three first letters from species). The tree includes the two newly sequenced orthologs from L’hoest’s monkey and Cotton-head tamarin (“new”). B) Results of the comprehensive positive selection analyses of primate (top row) and simian (bottom) TRIM69 sequences: BUSTED, MEME, FUBAR from HYPHY/Datamonkey.com, M1vsM2 and M7vsM8 from Bpp and PAML Codeml. Legend details: Size, length of the codon alignment; n. sp., number of species in the alignment; PS?, if the gene is under positive selection: Y, yes, N, no; p value, supporting a model under positive selection; PSS, positive selection sites; −, no site identified; omega (PS) in bpp, corresponds to the omega value in the positive selection class (dN/dS>1).

**Figure 6.**
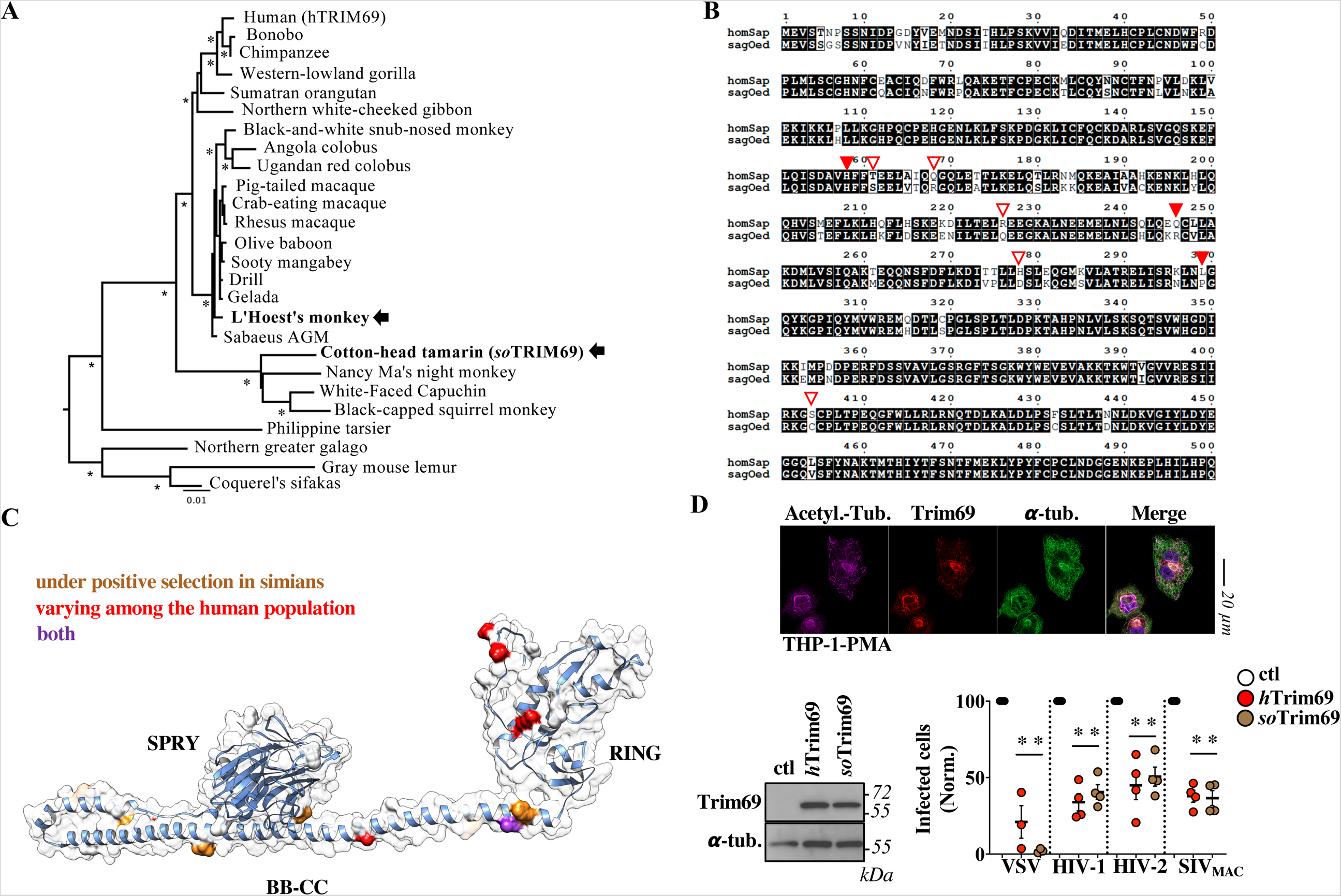
Trim69 exhibits signatures of positive selection during primate evolution and the divergent Trim69 orthologue from *Saguinus oedipus* (soTrim69) also inhibits viral infection and drives microtubule changes. A) Phylogenetic tree of primate Trim69, including two newly sequenced orthologs from L’hoest’s monkey and Cotton-head tamarin (arrows). PhyML was run with GTR+I+G model and 1,000 bootstrap replicates for node statistical supports (asterisks represent values above 700/1000). Sequence references are shown in **extended data Figure 11A**. B) Amino acid alignment of human and cotton-headed tamarin Trim69 (88.6% identity). Representation with ESPript [75]. Sites under positive selection are highlighted with red triangles (plain triangles, sites identified by >2 methods; open triangles, sites identified by one method). C) Amino acid positions undergoing positive selection or allelic variations in the human population are presented in the modeled 3D structure of human Trim69. D) Functional equivalency of the soTrim69 orthologue with respect to the induction of stable microtubule formation and to antiviral activities against VSV (n=3) and primate lentiviruses (n=4). Experiments present Avg and SEM. *, p<0.05 following a two-tailed Student t test between Trim69 proteins and respective controls.

