## Supplemental figures for "Trim69 is a microtubule regulator that acts as a pantropic viral inhibitor": Supplementary Figures Song et al. 2022.pdf

*Song et al., 2022, extended data Figure 1*

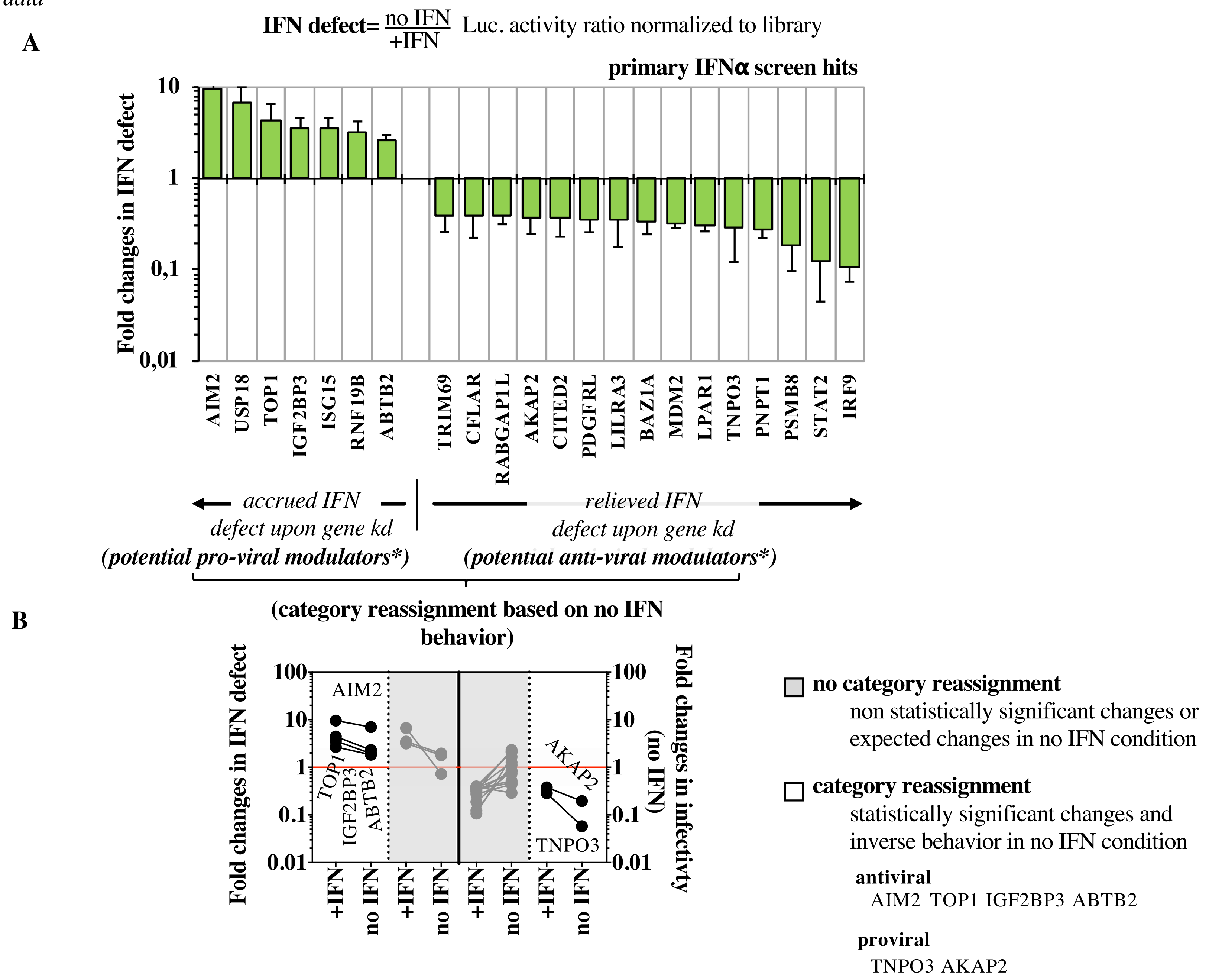

*Song et al., 2022, extended data*
*Figure 2*

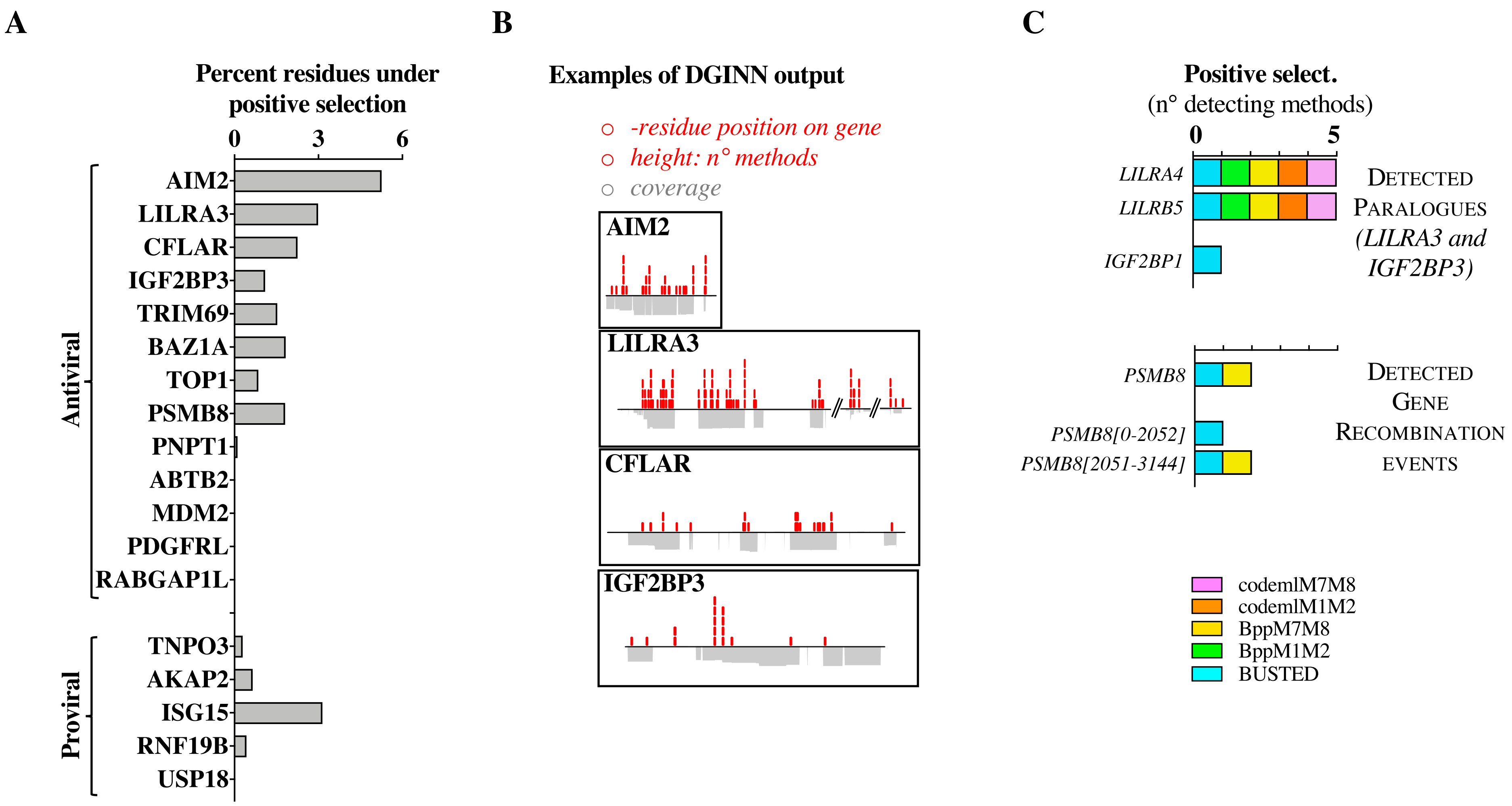

*Song et al., 2022, extended data*
*Figure 3*

**A**

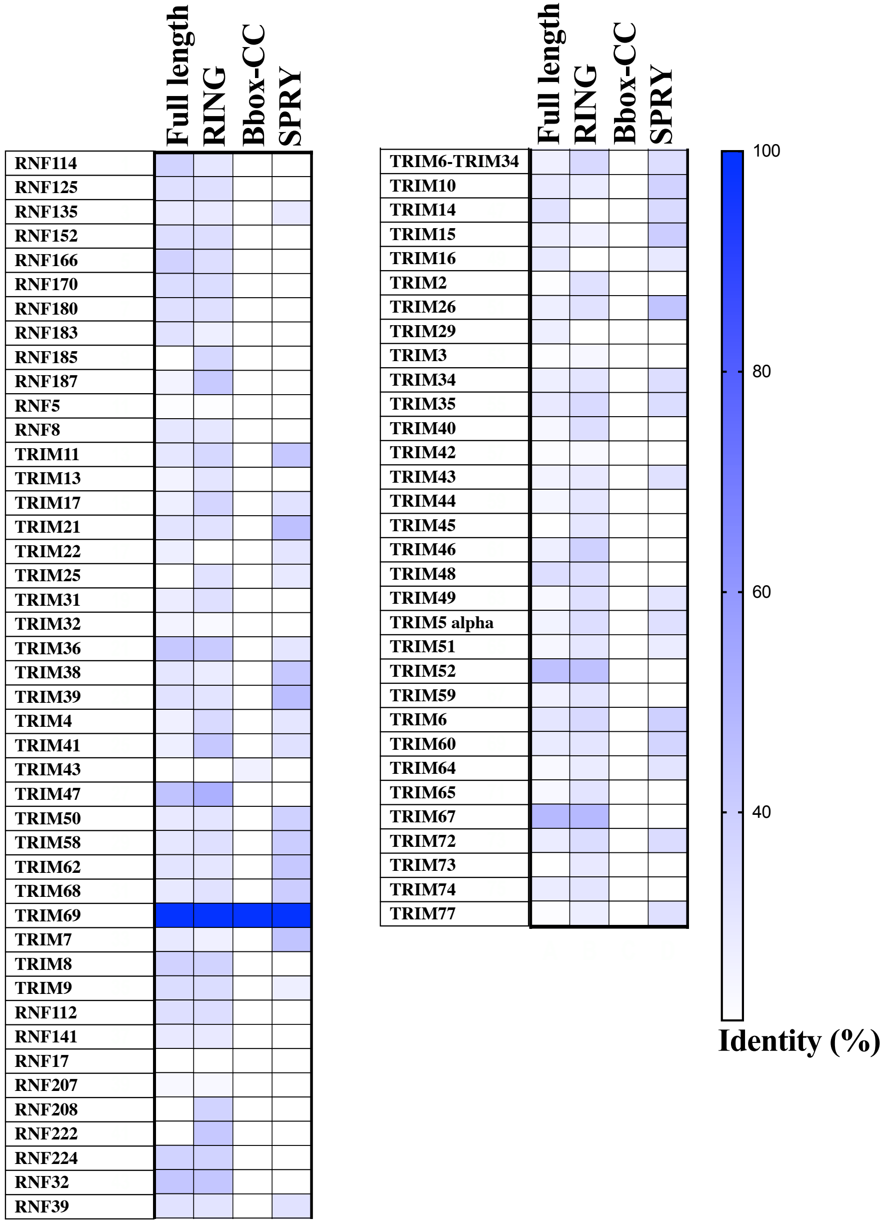


**B**

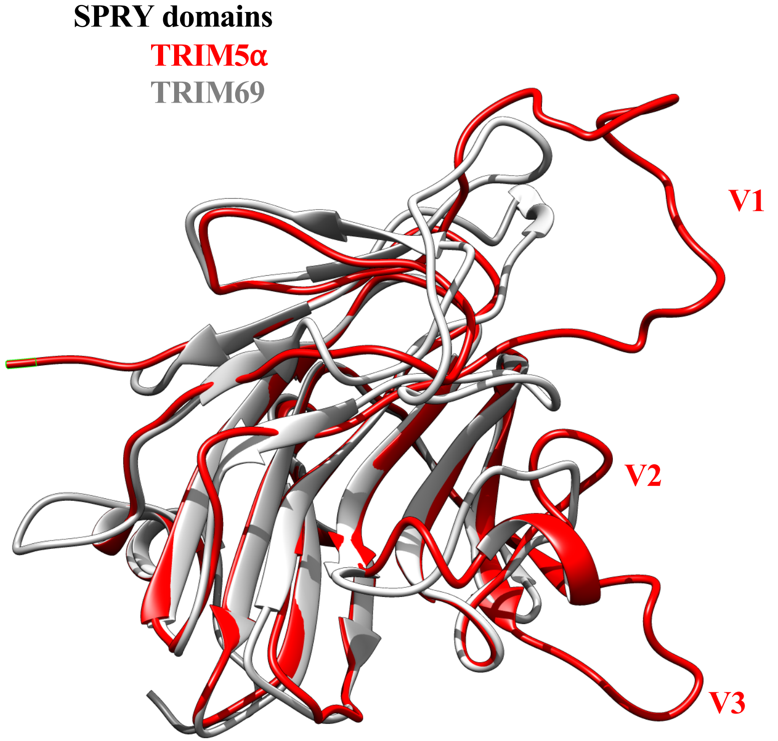

*Song et al., 2022, extended data*
*Figure 4*

**A**

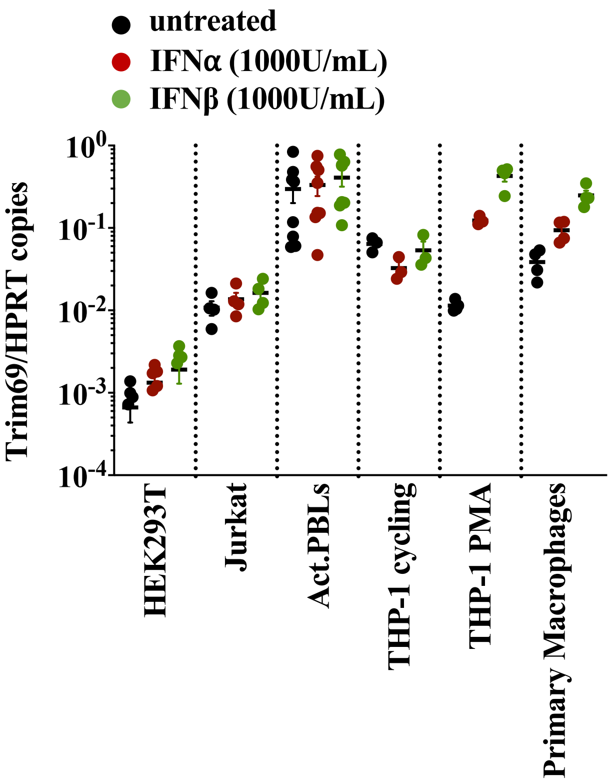


**B**

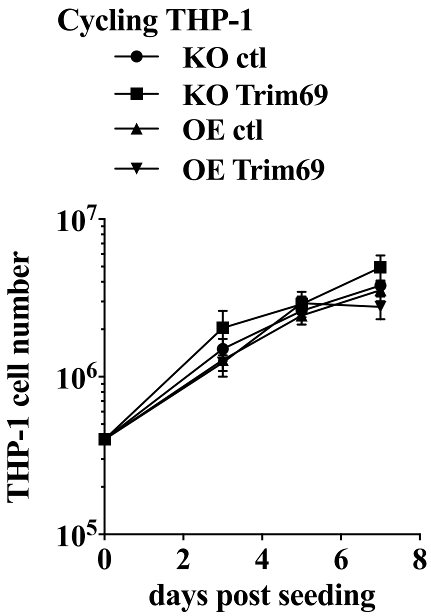

*Figure 5*

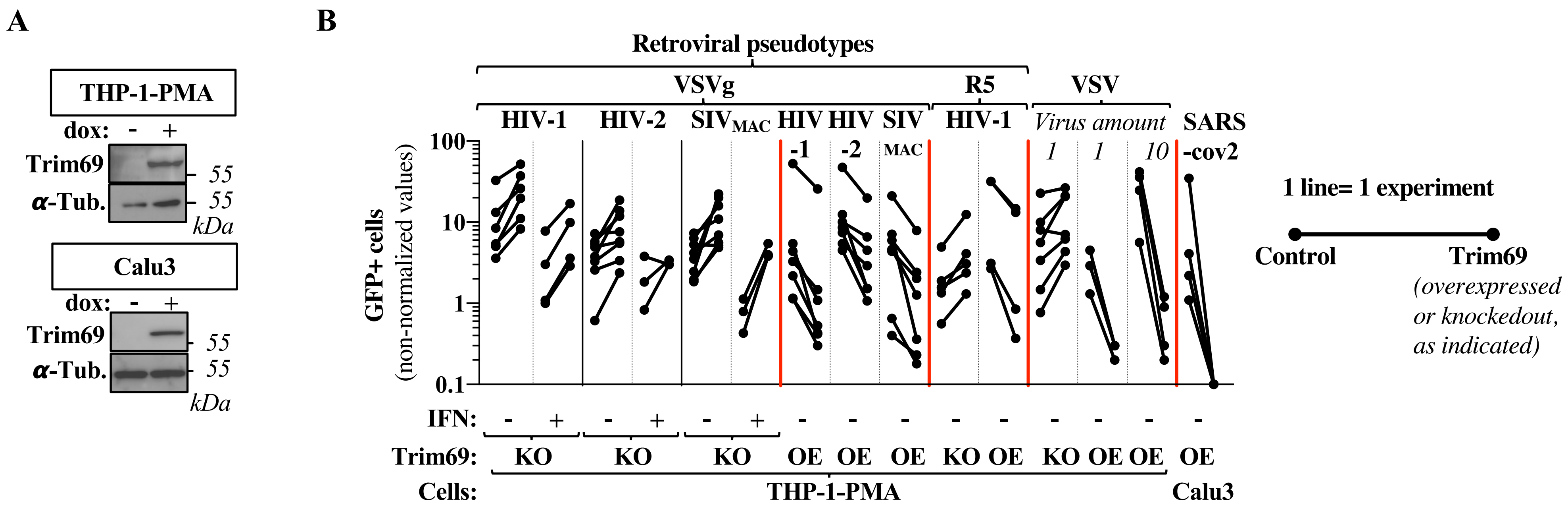
A
THP-1-PMA
dox: - +
Trim69
α-Tub.
55
55
kDa
Calu3
dox: - +
Trim69
α-Tub.
55
55
kDa
B
Retroviral pseudotypes
VSVg
R5
VSV
HIV-1
HIV-2
SIVMAC
HIV -1
HIV -2
SIV MAC
HIV-1
Virus amount
1
1
10
SARS -cov2
GFP+ cells (non-normalized values)
100
10
1
0.1
IFN: - + - + - + - - - - - - - - -
Trim69: KO KO KO OE OE OE KO OE KO OE OE OE
Cells: THP-1-PMA Calu3
1 line= 1 experiment
Control
Trim69 (overexpressed or knockedout, as indicated)

*Song et al., 2022, extended data*
*Figure 6*

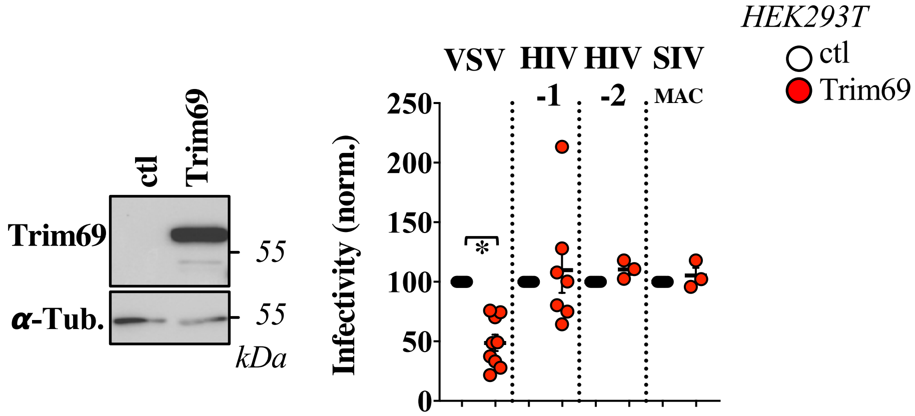

*Song et al., 2022, extended data*
*Figure 7*

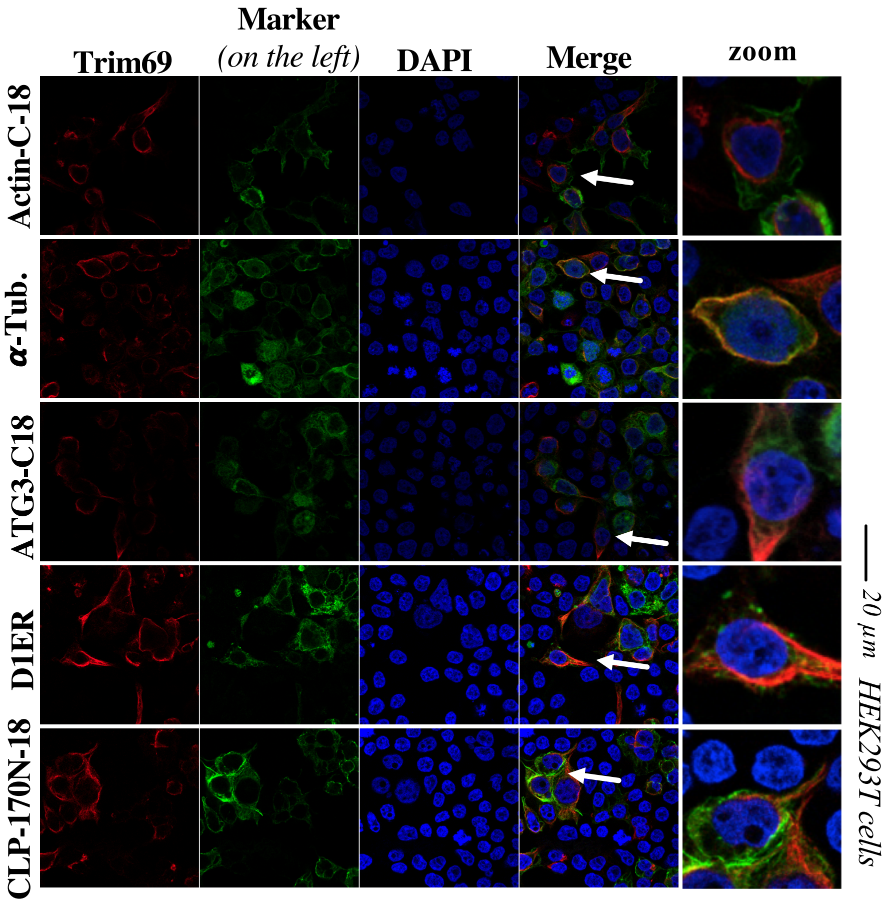


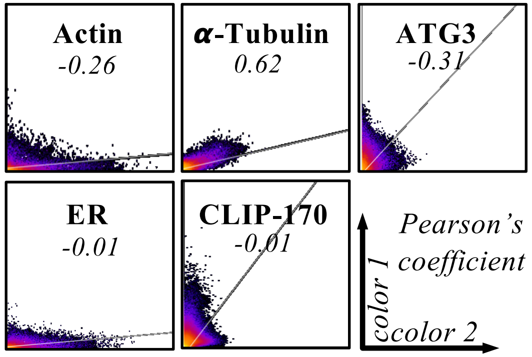

*Song et al., 2022, extended data*
*Figure 8*

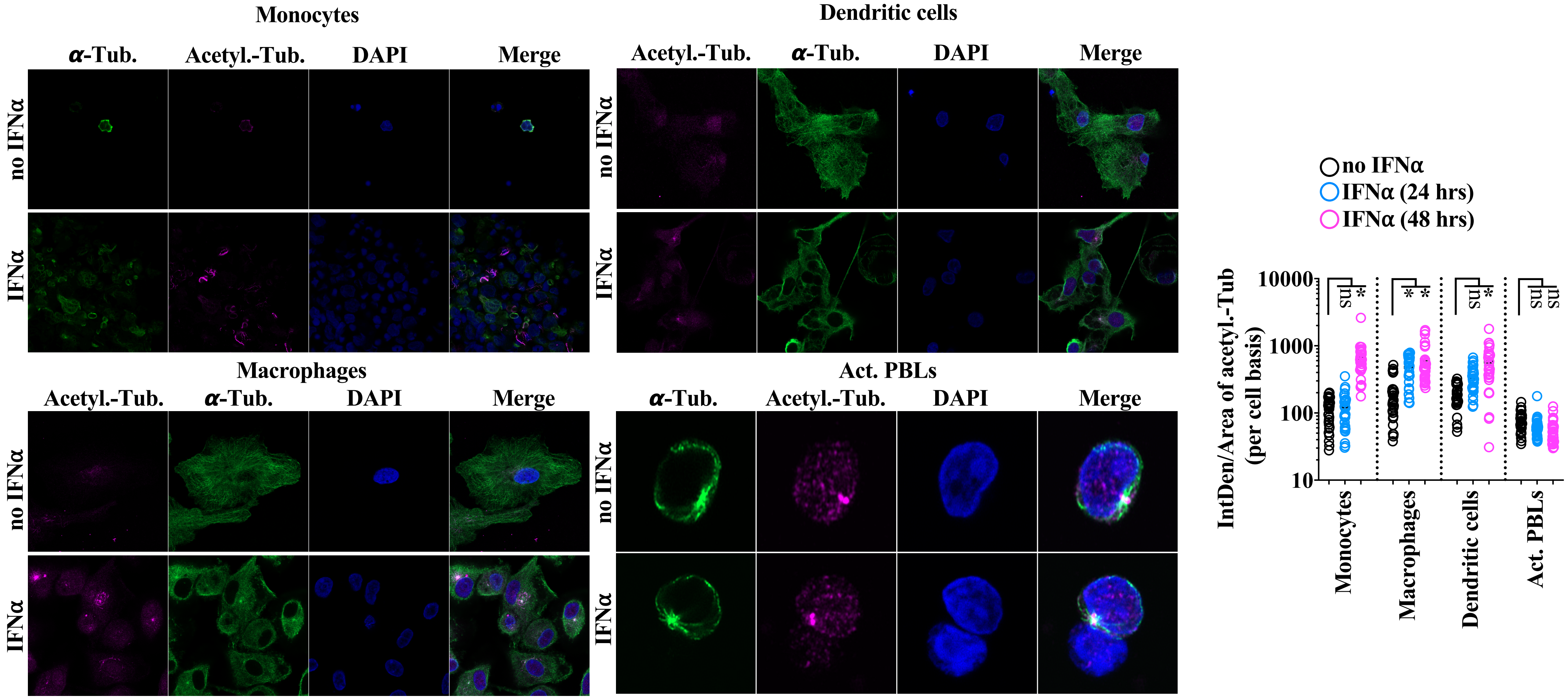

*Song et al., 2022, extended data*
*Figure 9*

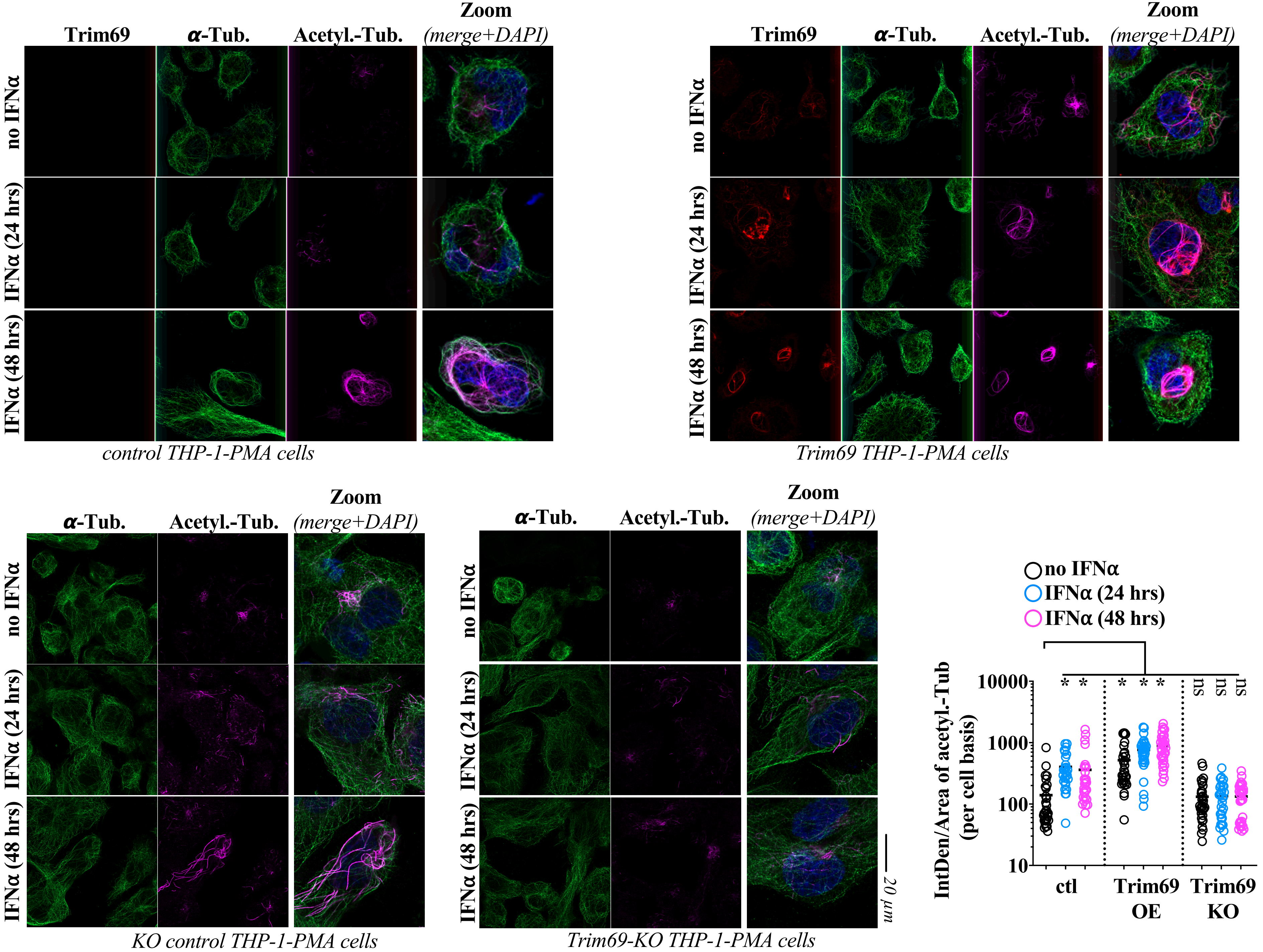

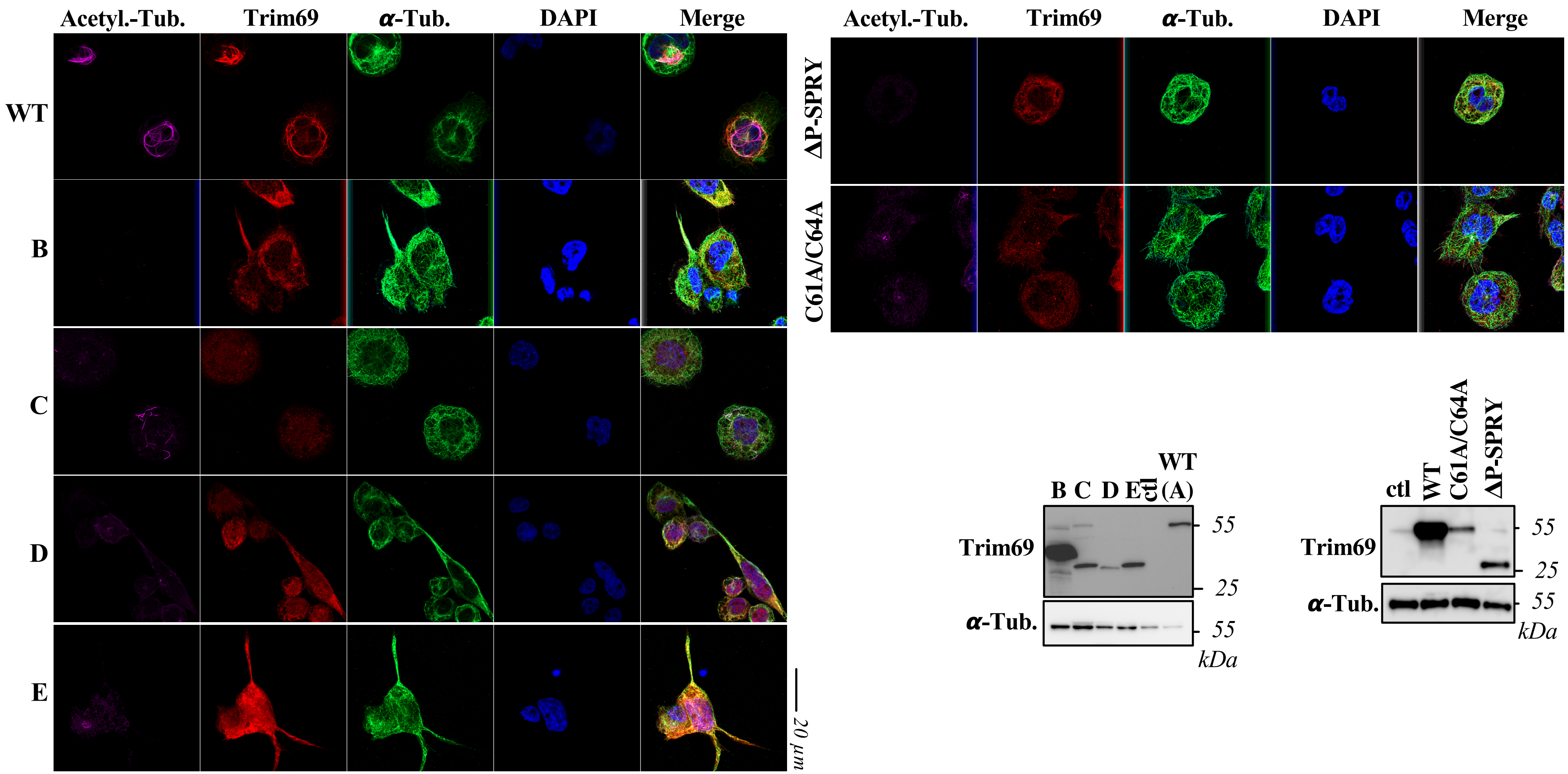
Acetyl.-Tub.
Trim69
α-Tub.
DAPI
Merge
WT
B
C
D
E
20 µm
Acetyl.-Tub.
Trim69
α-Tub.
DAPI
Merge
ΔP-SPRY
C61A/C64A
B
C
D
E
ctl
WT (A)
Trim69
α-Tub.
55
25
55
kDa
ctl
WT
C61A/C64A
ΔP-SPRY
Trim69
α-Tub.
55
25
55
kDa

**A**

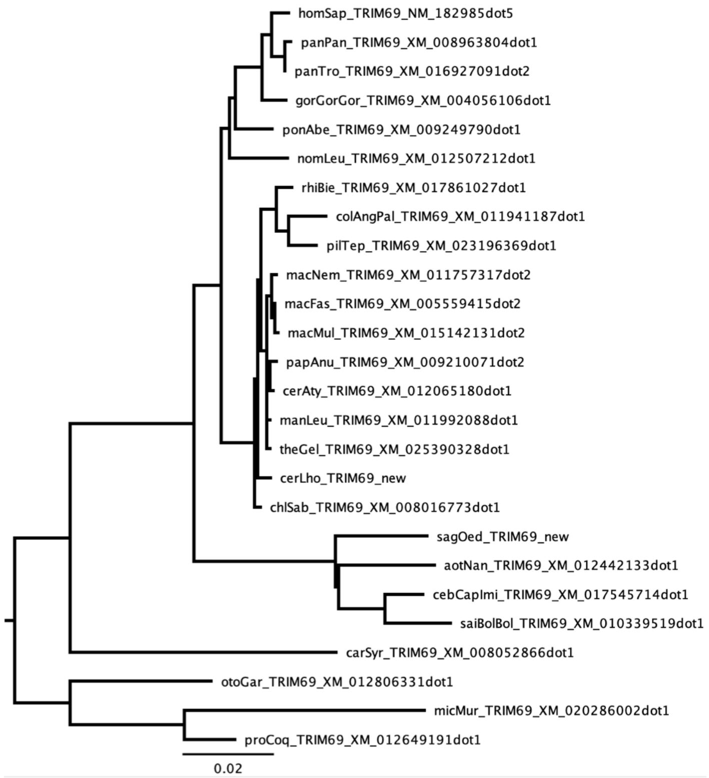


**B**

| Sequence alignment | | | | BUSTED | | MEME (p<0.1) | FUBAR (PP>0.9) | Bpp M1vsM2 | | | | Bpp M7vsM8 | | | | Codeml M1vsM2 | | | Codeml M7vsM8 | | |
|---|---|---|---|---|---|---|---|---|---|---|---|---|---|---|---|---|---|---|---|---|---|
| **Gene** | **Order** | **Size** | **n sp.** | **PS?** | **p value** | **PSS** | **PSS** | **PS?** | **p value** | **ω (PS)** | **PSS** | **PS?** | **p value** | **ω (PS)** | **PSS** | **PS?** | **p value** | **PSS** | **PS?** | **p value** | **PSS** |
| **TRIM69** | **primates** | 511 | 26 | **Y** | 0.0487 | 188, 274, 278, 288, 316, 332, 333, 364, 365, 437 | - | N | 0.1783 | 2.08 | **246** | N | 0.0796 | 3.12 | **246** | **Y** | 0,0119 | - | **Y** | 0.0116 | **246** |
| **TRIM69** | **simians** | 500 | 22 | N | 0.5 | **158, 246**, 278 | 161, 169, **158**, 226, **246** | N | 1 | 1.002 | - | N | 0.991 | 0 | - | **Y** | 0,0002 | **158, 246, 299** | **Y** | 0.0002 | **158, 246, 299,** 404 |
